# Neuronal PARIS-STAT3 axis drives tau pathology and glial activation in Alzheimer’s disease

**DOI:** 10.64898/2025.12.06.692741

**Authors:** Ji-Yang Song, Fatih Akkentli, Heejin Jo, Jinhee Park, Shinwon Ha, Javier Redding-Ochoa, Juan C. Troncoso, Sung-Ung Kang, Valina L. Dawson, Ted M. Dawson

## Abstract

PARIS, a substrate of Parkin, accumulates in Parkinson’s disease and promotes disease progression. Here, we demonstrate that PARIS also contributes to Alzheimer’s disease by elevating *STAT3* transcriptional activity, thereby inducing tau pathology, hippocampal atrophy, and glial activation. Genetic depletion of *Paris* reduced tau phosphorylation and cognitive decline in tauopathy mice, whereas neuron-specific PARIS overexpression caused tau accumulation, gliosis, and memory impairment. In contrast, astrocyte-specific overexpression did not induce pathology, indicating that PARIS acts in neurons to drive tau phosphorylation and glial activation. The pathological features induced by neuronal PARIS overexpression were rescued by STAT3 inhibition, demonstrating that PARIS–STAT3 signaling underlies these effects. Moreover, *Paris* knockout did not alter pathology in the amyloid-driven mouse model, highlighting specificity for tau pathology. Together, these findings reveal PARIS as a neuronal regulator of STAT3 signaling that exacerbates tau-mediated neurodegeneration and identify the PARIS–STAT3 pathway as a potential therapeutic target in Alzheimer’s disease.

## Introduction

Alzheimer’s disease (AD) is the most prevalent neurodegenerative disorder, yet effective disease-modifying therapies remain elusive^1,2^. Most therapeutic efforts have focused on reducing amyloid-β (Aβ) plaques, guided by the amyloid cascade hypothesis, supported by autosomal-dominant APP and presenilin mutations and by abundant Aβ deposition in AD brains^3–5^. This strategy led to the approval of antibody therapies that reduce amyloid plaque burden, yet cognitive benefits have been modest^6^. Consequently, focus has shifted toward tau pathology^7,8^. A notable case study showed that the homozygous APOE3 Christchurch (R136S) mutation delayed cognitive decline caused by the PSEN1-E280A mutation by nearly three decades^9,10^. This patient in her seventies exhibited mild tau pathology despite widespread amyloid deposition, compared with typical PSEN1-E280A mutation carriers who develop symptoms in their forties^9,10^. This case is consistent with the view that tau is a key determinant of cognitive impairment in AD^11,12^.

PARIS (ZNF746) was identified as a parkin-interacting substrate, and its pathogenic functions have been primarily investigated in the context of Parkinson’s disease (PD)^13–15^. In PD, inactivated parkin leads to accumulation of PARIS which represses the transcription of peroxisome proliferator-activated receptor gamma coactivator 1-α (PGC1α). Since PGC1α and its downstream genes are essential for mitochondrial function, reduced PGC1α contributes to neurodegeneration^15^. While these studies have established PARIS as a critical mediator of PD pathogenesis, accumulating evidence suggests that risk factors central to one neurodegenerative disorder can also contribute to the pathology of others^16–19^, and parkin, which is a key protein in PD, has been implicated in mitochondrial dysfunction in AD^20^. Based on this cross-disease principle, we examined whether PARIS also plays a role in AD. Our investigation revealed that PARIS contributes to AD by promoting tau pathology. Furthermore, using CUT&Tag^21^, a high-resolution chromatin profiling method, we identified *STAT3* as a critical transcriptional target of PARIS and uncovered a pathogenic mechanism by which PARIS contributes to tau-driven neurodegeneration in AD.

Our findings support a model in which neuronal perturbations precede glial alterations^22,23^, as neuron-specific PARIS overexpression was sufficient to induce STAT3 activation, tau pathology, and cognitive decline, whereas astrocyte-specific PARIS overexpression did not produce comparable effects. Taken together, these results not only identify a novel pathogenic mechanism of PARIS in AD but also highlight the PARIS–STAT3 axis as a therapeutic target.

## Results

### PARIS expression is increased in the brain tissue of Alzheimer’s disease patients

To determine whether PARIS expression is altered in the brain tissue of AD patients, we conducted analyses in regions implicated in AD pathology, including the hippocampus, entorhinal cortex, middle frontal gyrus, and superior/middle temporal gyrus (Fig. 1a–h)^24,25^. Compared with controls, PARIS levels were significantly increased in all four regions of AD brains. Total tau levels were unchanged, whereas phospho-tau markers AT8 (pSer202/pThr205) and AT270 (pThr181) were elevated, consistent with previous reports (Fig. 1a–h)^26,27^. By contrast, PARIS was not increased in the cerebellum, a region relatively unaffected in AD^28^, and AT8 and AT270 levels showed no difference from controls (Fig. 1i,j). These findings indicate a region-specific upregulation of PARIS in brain areas vulnerable to AD pathology, supporting a potential role for PARIS in the disease process.

**Fig. 1.**
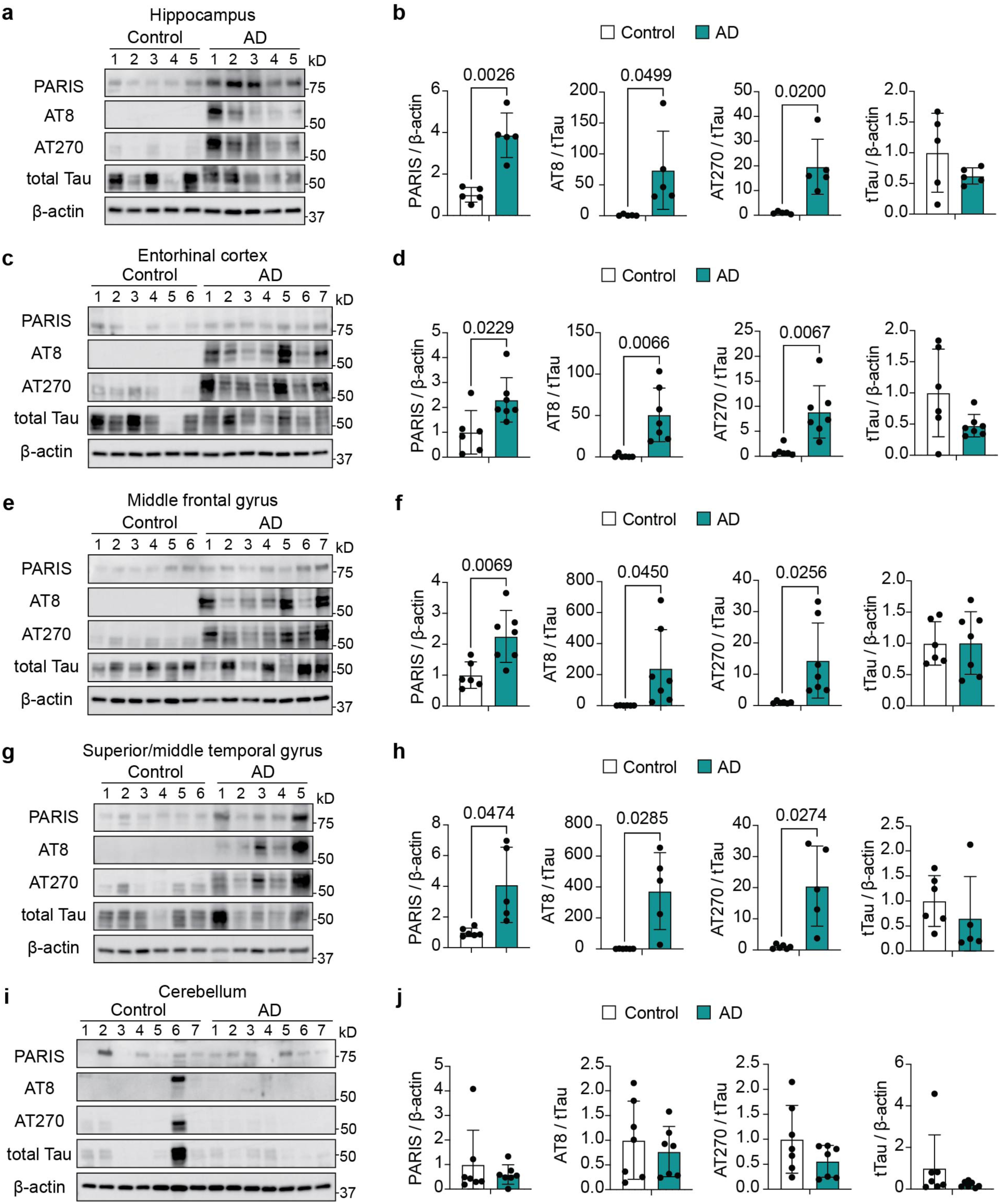
PARIS expression is increased in brain regions from AD patients. **a**, Immunoblot analysis of PARIS, phospho-tau (AT8 or AT270), total tau and β-actin in hippocampal samples from control and AD patients. **b**, Quantification of PARIS, phospho-tau and total tau levels from **a**. Data are mean ± SEM (Control, *n* = 5; AD, *n* = 5). Statistical significance was assessed by two-tailed *t*-test. **c**, Immunoblot analysis of PARIS, phospho-tau (AT8 or AT270), total tau and β-actin in entorhinal cortex from control and AD patients. **d**, Quantification of PARIS, phospho-tau and total tau levels from **c**. Data are mean ± SEM (Control, *n* = 6; AD, *n* = 7). Statistical significance was assessed by two-tailed *t*-test. **e**, Immunoblot analysis of PARIS, phospho-tau (AT8 or AT270), total tau and β-actin in middle frontal gyrus from control and AD patients. **f**, Quantification of PARIS, phospho-tau and total tau levels from **e**. Data are mean ± SEM (Control, *n* = 6; AD, *n* = 7). Statistical significance was assessed by two-tailed *t*-test. **g**, Immunoblot analysis of PARIS, phospho-tau (AT8 or AT270), total tau and β-actin in superior/middle temporal gyrus from control and AD patients. **h**, Quantification of PARIS, phospho-tau and total tau levels from **g**. Data are mean ± SEM (Control, *n* = 6; AD, *n* = 5). Statistical significance was assessed by two-tailed *t*-test. **i**, Immunoblot analysis of PARIS, phospho-tau (AT8 or AT270), total tau and β-actin in cerebellum from control and AD patients. **j**, Quantification of PARIS, phospho-tau and total tau levels from **i**. Data are mean ± SEM (Control, *n* = 7; AD, *n* = 7). Statistical significance was assessed by two-tailed *t*-test. Comparisons with *P* ≤ 0.05 are marked on the graph.

### Loss of *Paris* rescues behavioral deficits, tau phosphorylation, and neuronal integrity in PS19 mice

To investigate the effect of PARIS on tauopathy, we bred *Paris* knockout mice with PS19 mice, which carry the P301S mutant variant of the human microtubule-associated protein *MAPT*^29^. Behavioral analyses showed that deficits in PS19 mice were restored by *Paris* knockout (Fig. 2a–c). In the novel object recognition test^30^, PS19 mice showed a reduced discrimination index, whereas PS19;*Paris*^⁻/⁻^ mice performed similarly to WT. Total exploration time did not differ between groups (Fig. 2a). In the Y-maze novel-arm test^31^, the reduced time spent in the novel arm observed in PS19 mice was rescued by *Paris* deletion, while total distance and arm entries were unchanged (Fig. 2b). In cued fear conditioning^32^, baseline freezing before tone presentation did not differ among groups. However, after the tone, freezing responses increased in WT but remained lower in PS19 mice, indicating impaired associative fear memory. This deficit was ameliorated in PS19;*Paris*^⁻/⁻^ mice at each tone presentation (Fig. 2c).

**Fig. 2.**
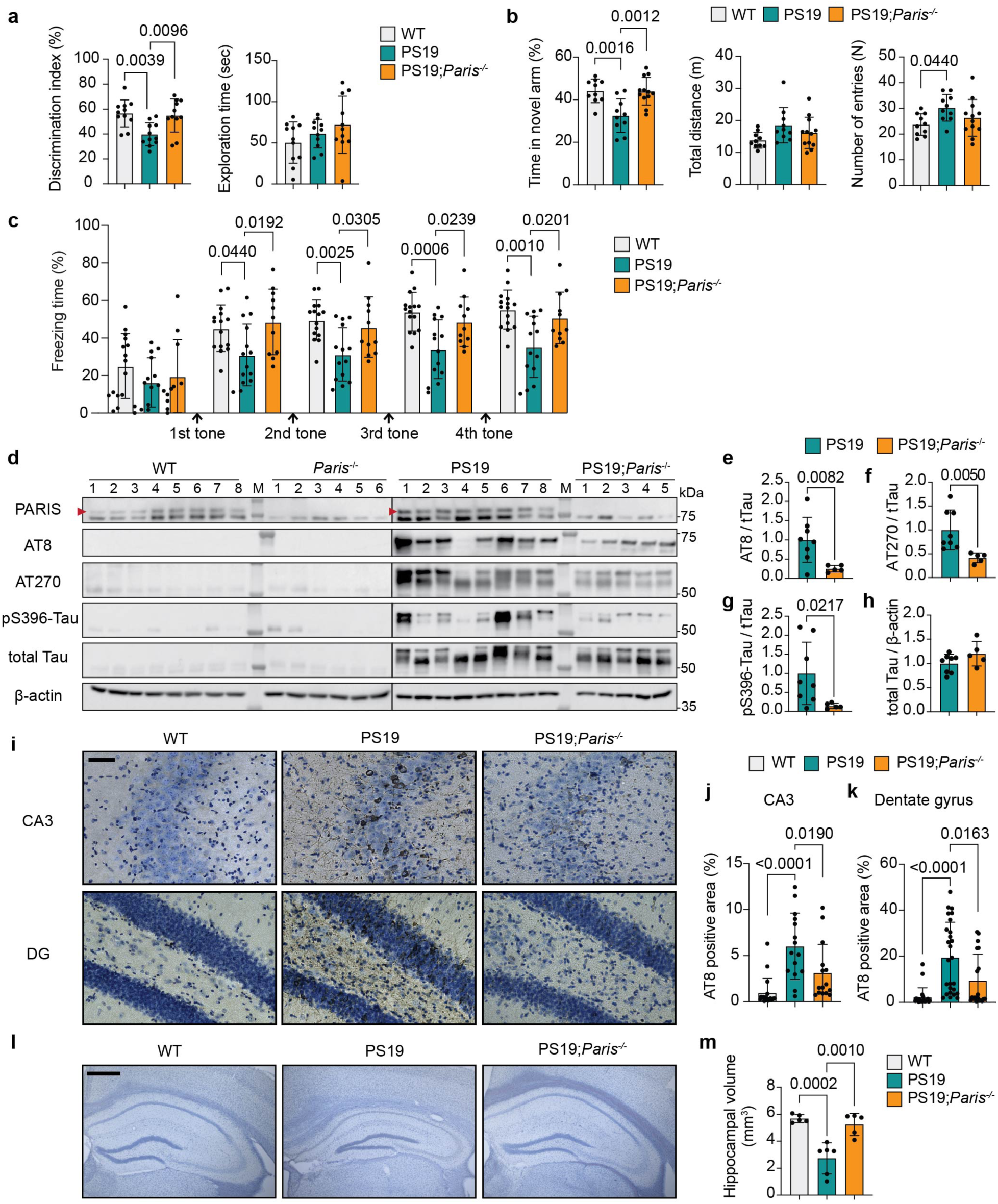
Loss of *Paris* rescues behavioral deficits, tau pathology, and hippocampal atrophy in PS19 mice. **a**, Percentage of time spent with the novel object and exploration time of the objects in the novel object recognition test. Data are mean ± SEM (WT, *n* = 11; PS19, *n* = 11; PS19;*Paris*^-/-^, *n* = 11). Statistical significance was assessed by one-way ANOVA followed by Tukey’s post hoc test. **b**, Percentage of time spent in the novel arm, total distance, and number of arm entries in the Y-maze test. Data are mean ± SEM (WT, *n* = 10; PS19, *n* = 10; PS19;*Paris*^-/-^, *n* = 12). Statistical significance was assessed by one-way ANOVA followed by Tukey’s post hoc test. **c**, Percentage of freezing time at baseline and after four stimulus tones in the fear conditioning test. Data are mean ± SEM (WT, *n* = 15; PS19, *n* = 14; PS19;*Paris*^-/-^, *n* = 11). Statistical significance was assessed by one-way ANOVA followed by Tukey’s post hoc test. **d**, Immunoblot of PARIS, pSer202/pThr205-tau (AT8), pThr181-tau (AT270), pSer396-tau, total tau and β-actin in the hippocampus of 10-month-old mice. The location of PARIS is indicated by arrows. **e**–**h**, Quantification of total tau, pSer202/pThr205-tau (AT8), pThr181-tau (AT270) and pSer396-tau from **d**. Data are mean ± SEM (PS19, *n* = 8; PS19;*Paris*^-/-^, *n* = 5). Statistical significance was assessed by two-tailed *t*-test. **i**, Representative IHC staining with the AT8 antibody in the CA3 region and dentate gyrus of the hippocampus in 10-month-old mice. Scale bar, 50 µm. **j**, Quantification of AT8-positive area in the CA3 from **i**. Data are mean ± SEM (WT, *n* = 19; PS19, *n* = 15; PS19;*Paris*^-/-^, *n* = 15). Statistical significance was assessed by one-way ANOVA followed by Tukey’s post hoc test (images from 3–5 sections per mouse, 4–6 mice per group). **k**, Quantification of AT8-positive area in the dentate gyrus from **i**. Data are mean ± SEM (WT, *n* = 22; PS19, *n* = 26; PS19;*Paris*^-/-^, *n* = 19). Statistical significance was assessed by one-way ANOVA followed by Tukey’s post hoc test (images from 3–7 sections per mouse, 4–5 mice per group). **l**, Representative images of Nissl-stained coronal sections of the hippocampus from 10-month-old mice. Scale bar, 500 µm. **m**, Quantification of hippocampal volume from **l**. Data are mean ± SEM (WT, *n* = 5; PS19, *n* = 6; PS19;*Paris*^-/-^, *n* = 5). Statistical significance was assessed by one-way ANOVA followed by Tukey’s post hoc test. Comparisons with *P* ≤ 0.05 are marked on the graph.

We next examined tau pathology in the hippocampus of 10-month-old littermates. Elevated levels of AT8, AT270, and pSer396-tau in PS19 mice were significantly reduced in PS19;*Paris*^⁻/⁻^ mice (Fig. 2d–h). In the entorhinal cortex, the absence of *Paris* decreased AT270 levels, while AT8 and pSer396-tau remained unchanged (Extended Data Fig. 1a–e). In the frontal cortex, AT270 and pSer396-tau were reduced by *Paris* deletion, whereas AT8 levels did not differ from PS19 (Extended Data Fig. 1f–j). These results indicate that *Paris* deletion leads to a reduction in tau phosphorylation in both hippocampal and cortical regions, with more pronounced effects in the hippocampus.

Consistent with these biochemical findings, tau aggregates in the CA3 and dentate gyrus of the hippocampus were also reduced in PS19;*Paris*^⁻/⁻^ mice (Fig. 2i–k).

We next quantified hippocampal volume to evaluate neuronal loss-associated structural changes. PS19 mice showed a marked reduction in hippocampal volume compared with WT controls, whereas *Paris* deletion significantly attenuated this loss (Fig. 2l,m). In parallel, immunostaining for the presynaptic marker SYNAPSIN I and the postsynaptic marker PSD95 revealed a pronounced decrease in synaptic area in the hippocampus of PS19 mice, which was restored in PS19;*Paris*^⁻/⁻^ mice (Extended Data Fig. 1k–m).

Together, these results indicate that loss of *Paris* mitigates not only tau pathology but also hippocampal atrophy and synaptic deficits in PS19 mice.

### Neuron-specific PARIS overexpression induces memory deficits and tau phosphorylation

To further investigate the impact of PARIS on tau pathology, we utilized transgenic mice that specifically overexpress human PARIS in neurons (CamK-PARIS)^13^. This model operates in a Tet-Off system, where PARIS expression was induced for 3 weeks by doxycycline withdrawal at 2 months of age. Neuron-specific PARIS-overexpressing mice exhibited reduced spatial memory and fear memory compared to littermate controls maintained on doxycycline-containing food, in which PARIS expression remained suppressed (Fig. 3a,b). In the Y-maze, CamK-PARIS mice spent less time in the novel arm compared to controls, despite no differences in total distance traveled or number of arm entries (Fig. 3a). In the cued fear conditioning test, freezing responses began to differ after the third tone, with neuronal PARIS-overexpressing mice exhibiting reduced freezing compared to controls (Fig. 3b).

**Fig. 3.**
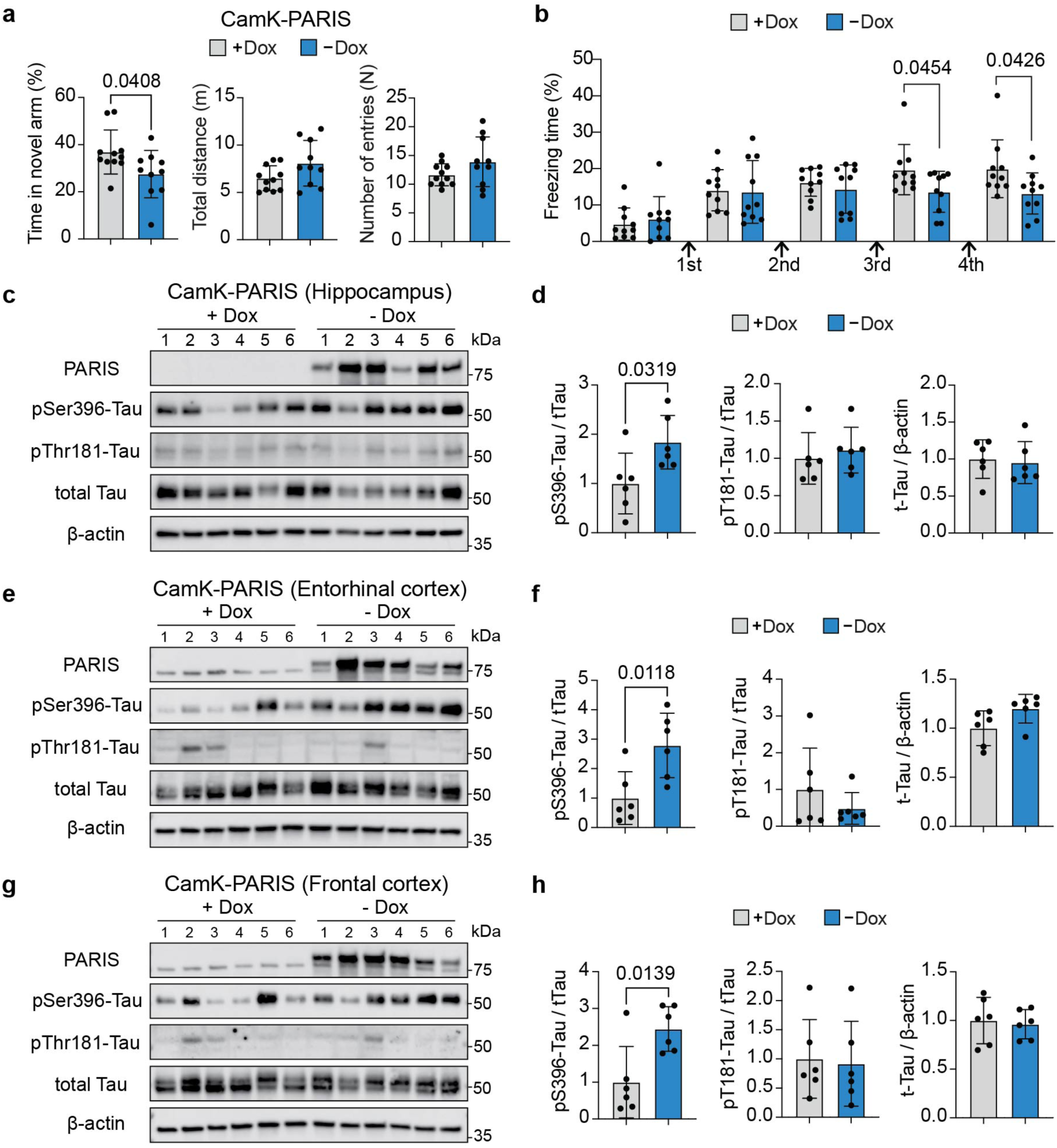
Neuron-specific PARIS overexpression causes memory loss and tau phosphorylation. PARIS expression was induced for 3 weeks by doxycycline withdrawal at 2 months of age. **a**, Percentage of time spent in the novel arm, total distance, and number of arm entries in the Y-maze test. Data are mean ± SEM (+Dox, *n* = 11; −Dox, *n* = 10). Statistical significance was assessed by two-tailed *t*-test. **b**, Percentage of freezing time at baseline and after four stimulus tones in the fear conditioning test with CamK-PARIS mice. Data are mean ± SEM (+Dox, *n* = 10; −Dox, *n* = 10). Statistical significance was assessed by two-tailed *t*-test. **c**, Immunoblot of PARIS, pSer396-tau, pThr181-tau (AT270), total tau and β-actin in the hippocampus of CamK-PARIS mice. PARIS expression was induced for 3 weeks by doxycycline withdrawal at 2 months of age. **d**, Quantification of pSer396-tau, pThr181-tau and total tau from **c**. Data are mean ± SEM (+Dox, *n* = 6; −Dox, *n* = 6). Statistical significance was assessed by two-tailed *t*-test. **e**, Immunoblot analysis of PARIS, pSer396-tau, pThr181-tau (AT270), total tau, and β-actin in entorhinal cortex samples from CamK-PARIS mice. **f**, Quantification of pSer396-tau, pThr181-tau, and total tau levels from **e**. Data are mean ± SEM (+Dox, *n* = 6; −Dox, *n* = 6). Statistical significance was assessed by two-tailed *t*-test. **g**, Immunoblot analysis of PARIS, pSer396-tau, pThr181-tau (AT270), total tau, and β-actin in frontal cortex samples from CamK-PARIS mice. **h**, Quantification of pSer396-tau, pThr181-tau, and total tau levels from **g**. Data are mean ± SEM (+Dox, *n* = 6; −Dox, *n* = 6). Statistical significance was assessed by two-tailed *t*-test. Comparisons with *P* ≤ 0.05 are marked on the graph.

In the hippocampus of these mice, PARIS overexpression increased tau phosphorylation at Ser396, whereas phosphorylation at Thr181 remained unchanged (Fig. 3c,d). Similar results were observed in the entorhinal and frontal cortices, where pSer396-tau was elevated without a corresponding change in pThr181-tau (Fig. 3e–h). Collectively, these analyses showed residue-specific changes, while total tau levels remained unchanged across all examined regions (Fig. 3a–h). This selective effect may reflect the relatively short duration of PARIS induction (3 weeks), which was insufficient to drive phosphorylation at additional sites. Extending PARIS induction was not feasible, as CamK-PARIS mice exhibited a rapid decline in survival one month after switching to a regular diet^13^.

### PARIS acts as a transcriptional activator of *STAT3*

The next objective was to investigate the mechanism through which PARIS regulates tau phosphorylation and cognitive impairment in mice. PGC1α had been identified as a target gene for PARIS, and its mechanism had been extensively investigated in PD^13–15,33^ Therefore, we checked the levels of PGC1α to determine if the same mechanism applied to tau pathology. However, there was no change in PGC1α expression in *Paris* knockout PS19 mice (Extended Data Fig. 2a,b).

To identify new transcriptional targets of PARIS relevant to tau pathology, we performed CUT&Tag in hippocampal tissue from a mouse with neuron-specific overexpression of FLAG-tagged PARIS. CUT&Tag is a next-generation chromatin profiling approach that provides higher sensitivity and lower background than conventional ChIP-seq, thereby allowing reliable mapping of protein–DNA interactions even from limited material^21,34,35^. PARIS expression in the hippocampal samples was confirmed by PARIS and FLAG immunoblotting (Extended Data Fig. 2c). We obtained 20.9 million and 56.2 million mapped reads for PARIS-FLAG and IgG controls, respectively, and identified 99K peaks in the PARIS-FLAG sample normalized to IgG (FRiP = 0.58)^36^, with heatmaps (±3 kb) showing enrichment in PARIS-FLAG but not IgG (Fig. 4a and Extended Data Fig. 2d).

**Fig. 4.**
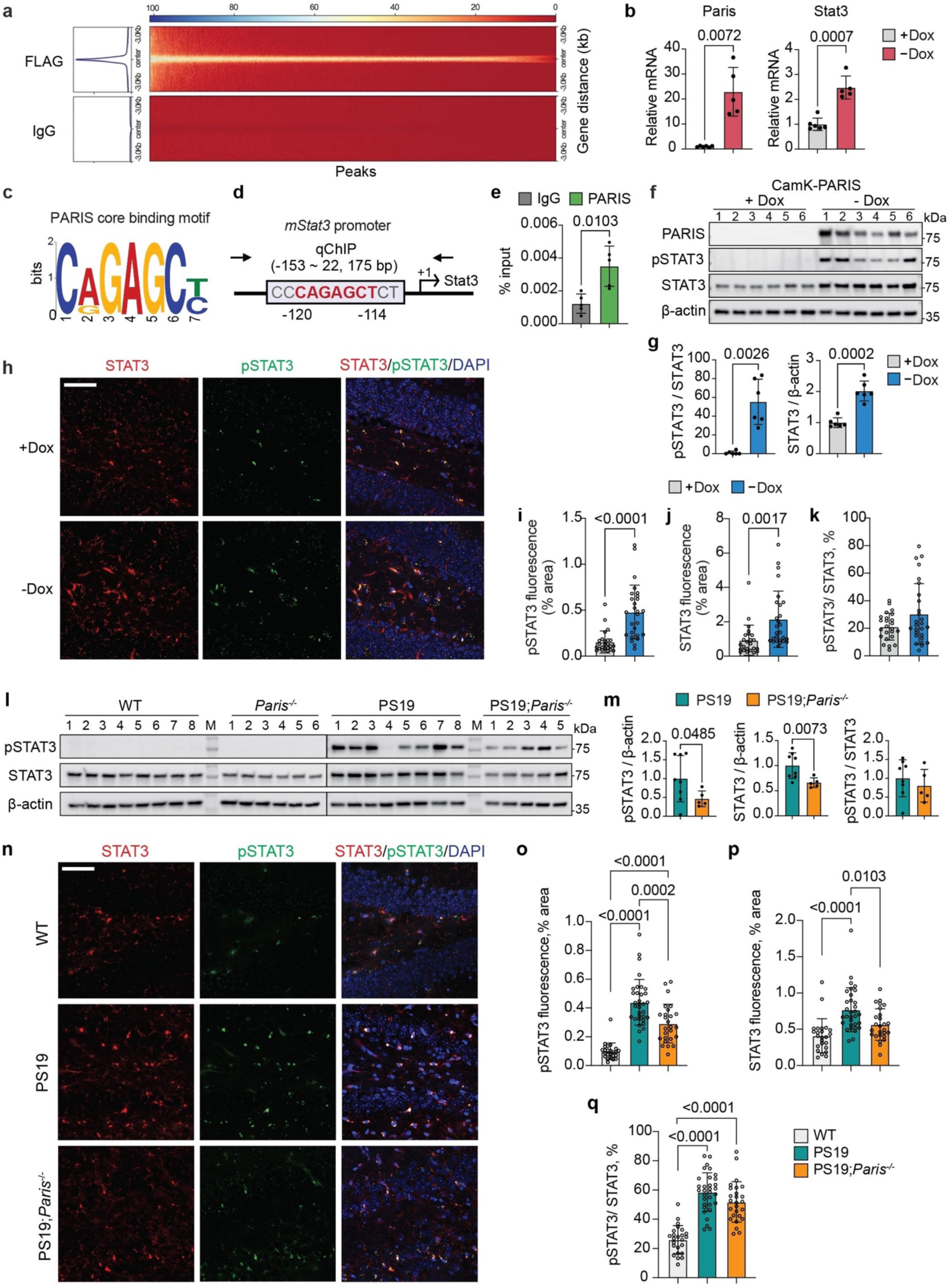
Neuron-specific PARIS overexpression increases *Stat3* transcription and activation, which are rescued by *Paris* knockout in PS19 mice. **a,** Heat map of CUT&Tag peaks. **b**, Relative mRNA expression of Paris and Stat3 normalized to β-actin by quantitative RT–PCR. Data are mean ± SEM (+Dox, *n* = 6; −Dox, *n* = 15). Statistical significance was assessed by two-tailed *t*-test. **c**, Core binding sequence of PARIS obtained by CUT&Tag analysis. **d**, Location of primers for quantitative ChIP assay (arrows) in the mouse *Stat3* promoter construct. **e**, Quantitative ChIP assay showing enrichment of PARIS at the Stat3 promoter in the hippocampus of CamK-PARIS mice. Data are mean ± SEM (+Dox, *n* = 5; −Dox, *n* = 5). Statistical significance was assessed by two-tailed *t*-test. **f**, Immunoblot analysis of PARIS, pTyr705-STAT3, total STAT3, and β-actin in hippocampal samples from CamK-PARIS mice. PARIS expression was induced for 3 weeks by doxycycline withdrawal starting at 2 months of age. **g**, Quantification of pTyr705-STAT3 and total STAT3 levels from **f**. Data are mean ± SEM (+Dox, *n* = 6; −Dox, *n* = 6). Statistical significance was assessed by two-tailed *t*-test. **h**, Representative co-immunostaining images of STAT3 (red), pTyr705-STAT3 (green), and DAPI (blue) in the hippocampus of CamK-PARIS mice. Scale bar, 50 µm. **i–k**, Quantification of STAT3-positive area, pTyr705-STAT3–positive area, and the ratio of pTyr705-STAT3 to STAT3 from **h**. Data are mean ± SEM (+Dox, *n* = 24; −Dox, *n* = 27; images from 4–6 sections per mouse, 5 mice per group). Statistical significance was assessed by two-tailed *t*-test. **l**, Immunoblot analysis of pTyr705-STAT3, total STAT3, and β-actin in hippocampal samples from 10-month-old PS19 mice. The same mouse samples as in Fig. 2d were used. **m**, Quantification of total STAT3 levels, pTyr705-STAT3 levels, and the ratio of pTyr705-STAT3 to STAT3 from **l**. Data are mean ± SEM (PS19, *n* = 8; PS19;*Paris*^-/-^, *n* = 5). Statistical significance was assessed by two-tailed *t*-test. **n**, Representative co-immunostaining images of STAT3 (red), pTyr705-STAT3 (green), and DAPI (blue) in the hippocampus of 10-month-old PS19 mice. Scale bar, 50 µm. **o–q**, Quantification of STAT3-positive area, pTyr705-STAT3–positive area, and the ratio of pTyr705-STAT3 to STAT3 from **n**. Data are mean ± SEM (WT, *n* = 24; PS19, *n* = 32; PS19;*Paris*^-/-^, *n* = 26; images from 4–6 sections per mouse and 5–6 mice per group). Statistical significance was assessed by one-way ANOVA followed by Tukey’s post hoc test. Comparisons with *P* ≤ 0.05 are marked on the graph.

Of the 99K peaks, 24.6% localized to promoter regions, corresponding to 1,139 annotated genes. Pathway analysis identified 91 genes across 10 significant pathways, with *Stat3* and *Mapk8* among the most frequently represented (Extended Data Fig. 2e). Among these, Stat3 mRNA was increased in hippocampi of PARIS-overexpressing mice (Fig. 4b), whereas Mapk8 mRNA remained unchanged (Extended Data Fig. 2f). STAT3-related pathways are listed in Extended Data Fig. 2g. Motif analysis of CUT&Tag peaks identified a consensus PARIS binding sequence (E-value = 7.5 × 10⁻⁹; Fig. 4c), and this motif was present within the *Stat3* promoter region (Fig. 4d), providing mechanistic support for direct transcriptional regulation of *Stat3* by PARIS. Chromatin immunoprecipitation further validated PARIS occupancy at the endogenous *Stat3* promoter (Fig. 4e and Extended Data Fig. 3h).

PARIS overexpression increased both total STAT3 and pTyr705-STAT3 in CamK-PARIS mice (Fig. 4f,g). Immunofluorescence confirmed these findings, showing elevated levels of total STAT3 and pTyr705-STAT3 consistent with the immunoblot results (Fig. 4h–j), although the ratio of pTyr705-STAT3 to total STAT3 remained unchanged (Fig. 4k).

These findings suggest that the increase in phosphorylated STAT3 can be largely attributable to the overall elevation in total STAT3 rather than a change in phosphorylation efficiency. The lack of detectable endogenous pTyr705-STAT3 in control immunoblots may be explained by low basal abundance and/or instability under RIPA extraction conditions (Fig. 4f). To assess whether PARIS is required for STAT3 regulation *in vivo*, we examined PS19 mice lacking *Paris*. In the hippocampus of PS19 mice, both total STAT3 and pTyr705-STAT3 were reduced by *Paris* knockout (Fig. 4l,m), whereas the relative proportion of pTyr705-STAT3 to total STAT3 remained unchanged (Fig. 4m). This suggests that PARIS directly regulates STAT3 expression but does not independently affect STAT3 phosphorylation. Immunofluorescence analysis validated these findings (Fig. 4n–q).

### PARIS induces astrogliosis and microglial activation

STAT3 has been established as a critical regulator of astrogliosis, primarily in astrocytes, and its close association with AD pathology has been demonstrated in previous studies^37–40^. Given that neuronal PARIS overexpression increases STAT3 expression, we hypothesized that PARIS may influence behavioral deficits and tau pathology in mice by promoting astrogliosis through neuronal STAT3 signaling.

Consistent with this possibility, the elevated GFAP expression in PS19 mice was reduced by *Paris* deletion, whereas IBA1 levels remained unchanged (Fig. 5a,b). To further assess astrocytic reactivity, we performed double immunostaining for GFAP and complement C3, a marker of neurotoxic reactive astrocytes^41^. Both GFAP and C3 intensities were increased in PS19 mice and reduced by *Paris* knockout (Fig. 5c–e).

**Fig. 5.**
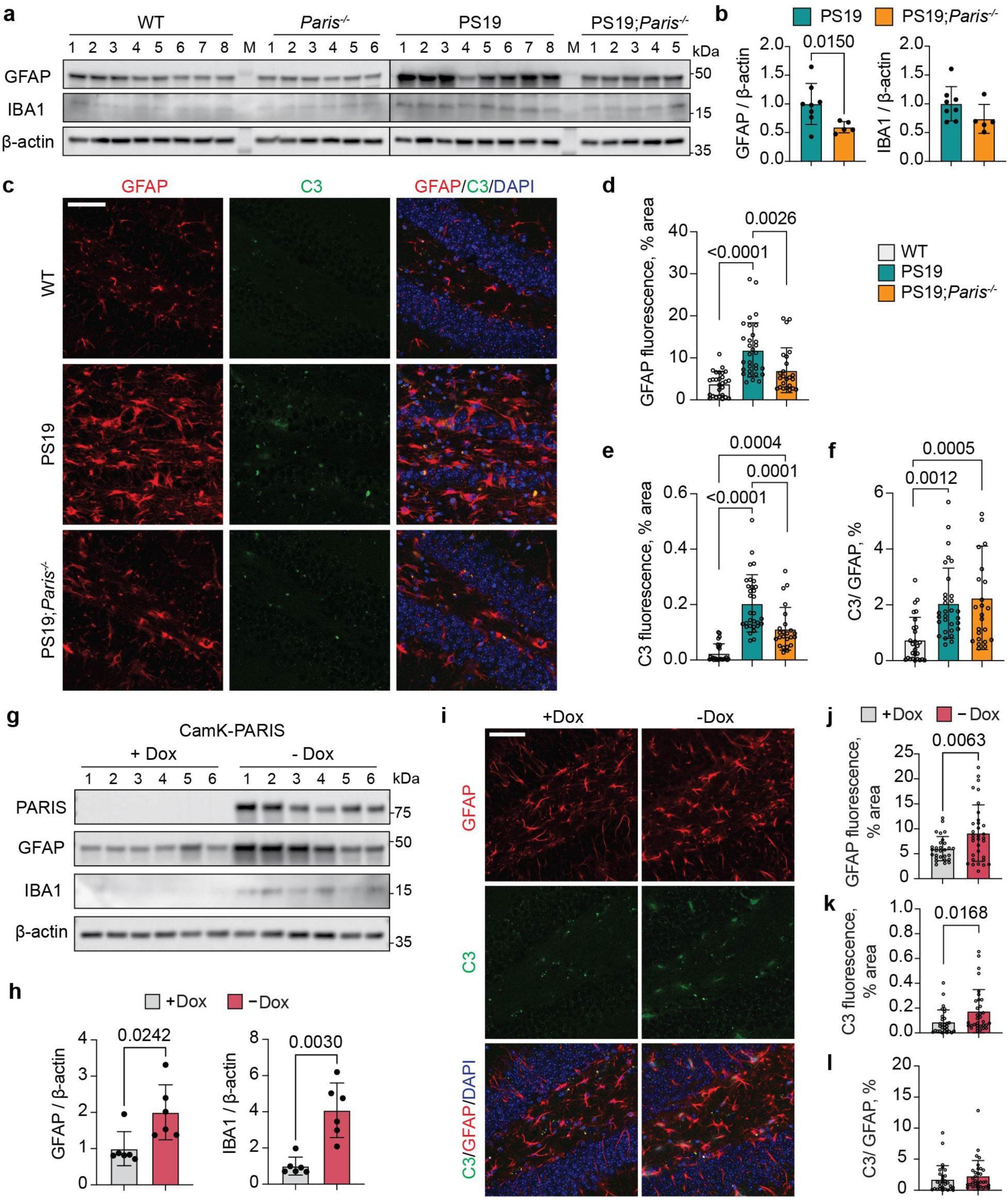
Neuron-specific PARIS overexpression enhances astrocytic activation, which is alleviated by *Paris* knockout in PS19 mice. **a,** Immunoblot analysis of GFAP, IBA1, and β-actin in hippocampal samples from 10-month-old PS19 mice. The same mouse samples as in Fig. 2d were used. **b**, Quantification of GFAP and IBA1 levels from **a**. Data are mean ± SEM (PS19, *n* = 8; PS19;*Paris*^-/-^, *n* = 5). Statistical significance was assessed by two-tailed *t*-test. **c**, Representative co-immunostaining images of GFAP (red), C3 (green), and DAPI (blue) in the hippocampus of 10-month-old PS19 mice. Scale bar, 50 µm. **d–f**, Quantification of GFAP, C3, and relative C3 fluorescence from **c**. Data are mean ± SEM (WT, *n* = 27; PS19, *n* = 31; PS19;*Paris*^-/-^, *n* = 24; images from 4–6 sections per mouse and 5–6 mice per group). Statistical significance was assessed by one-way ANOVA followed by Tukey’s post hoc test. **g**, Immunoblot analysis of GFAP, IBA1, and β-actin in hippocampal samples from CamK-PARIS mice. PARIS expression was induced for 3 weeks by doxycycline withdrawal starting at 2 months of age. **h**, Quantification of GFAP and IBA1 levels from **g**. Data are mean ± SEM (+Dox, *n* = 6; −Dox, *n* = 6). Statistical significance was assessed by two-tailed *t*-test. **i**, Representative co-immunostaining images of GFAP (red), C3 (green), and DAPI (blue) in the hippocampus of CamK-PARIS mice. Scale bar, 50 µm. **j–l**, Quantification of GFAP, C3 and relative C3 fluorescence from **i**. Data are mean ± SEM (+Dox, *n* = 29; −Dox, *n* = 33; images from 5–8 sections per mouse and 5 mice per group). Statistical significance was assessed by two-tailed *t*-test. Comparisons with *P* ≤ 0.05 are marked on the graph.

However, the ratio of C3 to GFAP remained unchanged (Fig. 5f), indicating that PARIS increases astrocyte number and overall reactivity, but does not further increase the activation state of individual astrocytes.

Microglial responses were examined by co-labeling IBA1 with CD16/CD32, a marker of activated microglia^42^. Consistent with the immunoblot results, IBA1 intensity was not affected by *Paris* deletion, but the ratio of CD16/CD32 to IBA1 was decreased in PS19;*Paris*^⁻/⁻^ mice (Extended Data Fig. 3a,b). These findings suggest that PARIS does not alter microglial numbers but enhances their activation.

To further assess the impact of neuronal PARIS overexpression on gliosis, we analyzed CamK-PARIS mice. GFAP levels were increased in these mice (Fig. 5g,h), consistent with the reduction observed in PS19;*Paris*^⁻/⁻^ mice, indicating that both models point to a similar role of PARIS in astrocytic regulation. Immunostaining with GFAP and C3 confirmed this effect, showing elevated astrocytic reactivity, while the C3-to-GFAP ratio remained unchanged (Fig. 5i–l). Thus, neuronal PARIS promotes astrocytic expansion and reactivity consistent with the knockout findings. For microglia, neuronal PARIS overexpression increased IBA1 immunoreactivity (Fig. 5g,h). In addition, both IBA1 intensity and the ratio of CD16/CD32 to IBA1, demonstrating that PARIS enhances not only microglial abundance but also their activation (Extended Data Fig. 3c,d). This parallels the knockout model, where Paris deletion reduced microglial activation. The difference in IBA1 expression between the two models may reflect the stronger and more rapid effects of neuronal PARIS overexpression, which is severe enough to cause premature mortality in CamK-PARIS mice within weeks. Collectively, these results reinforce the role of neuronal PARIS as a potent driver of both astrocytic and microglial responses.

### Astrocyte-specific PARIS overexpression does not induce behavioral deficits and tau pathology

Given that STAT3 function has been primarily emphasized in astrocytes, and that both neuronal PARIS overexpression and *Paris* deletion in tauopathy models consistently demonstrated PARIS-dependent increases in astrocytic markers, we next investigated the effects of astrocyte-specific PARIS overexpression. We generated GFAP-PARIS mice by crossing *GFAP*-*tTA* and *tet*-*PARIS* lines, applying the same Tet-Off system used in CamK-PARIS mice. At 3 weeks of PARIS induction, no differences were observed in Y-maze performance (data not shown). Behavioral testing was therefore extended to 6 weeks, at which point both Y-maze and cued fear conditioning were conducted. Despite a twofold longer induction period compared to CamK-PARIS mice, astrocyte-specific PARIS overexpression did not affect spatial and fear memory (Extended Data Fig. 4a,b).

Biochemical analyses of the hippocampus, entorhinal cortex, and frontal cortex revealed no increase in tau phosphorylation (Extended Data Fig. 4c–h). In contrast, both total STAT3 and phosphorylated STAT3, as well as GFAP expression, were markedly elevated across all examined regions, confirming robust astrocytic reactivity. IBA1 levels were unchanged in the hippocampus and frontal cortex, but were dramatically elevated in the entorhinal cortex (Extended Data Fig. 4c–h).Together, these results demonstrate that while PARIS is a potent regulator of STAT3 expression, activation, and astrocytic expansion, its neuronal expression is critical for driving tau phosphorylation and cognitive decline.

### STAT3 inhibition alleviates PARIS-driven behavioral deficits and neuropathological phenotypes

To test whether STAT3 mediates the effects of neuronal PARIS, we treated CamK-PARIS mice with the STAT3 inhibitor napabucasin (20 mg/kg, orally, daily) starting at the time of doxycycline withdrawal (Fig. 6a)^43,44^. Control littermates carrying only one transgene (*CamK*-*tTA* or *tet*-*PARIS*) were maintained under the same dietary conditions.

**Fig. 6.**
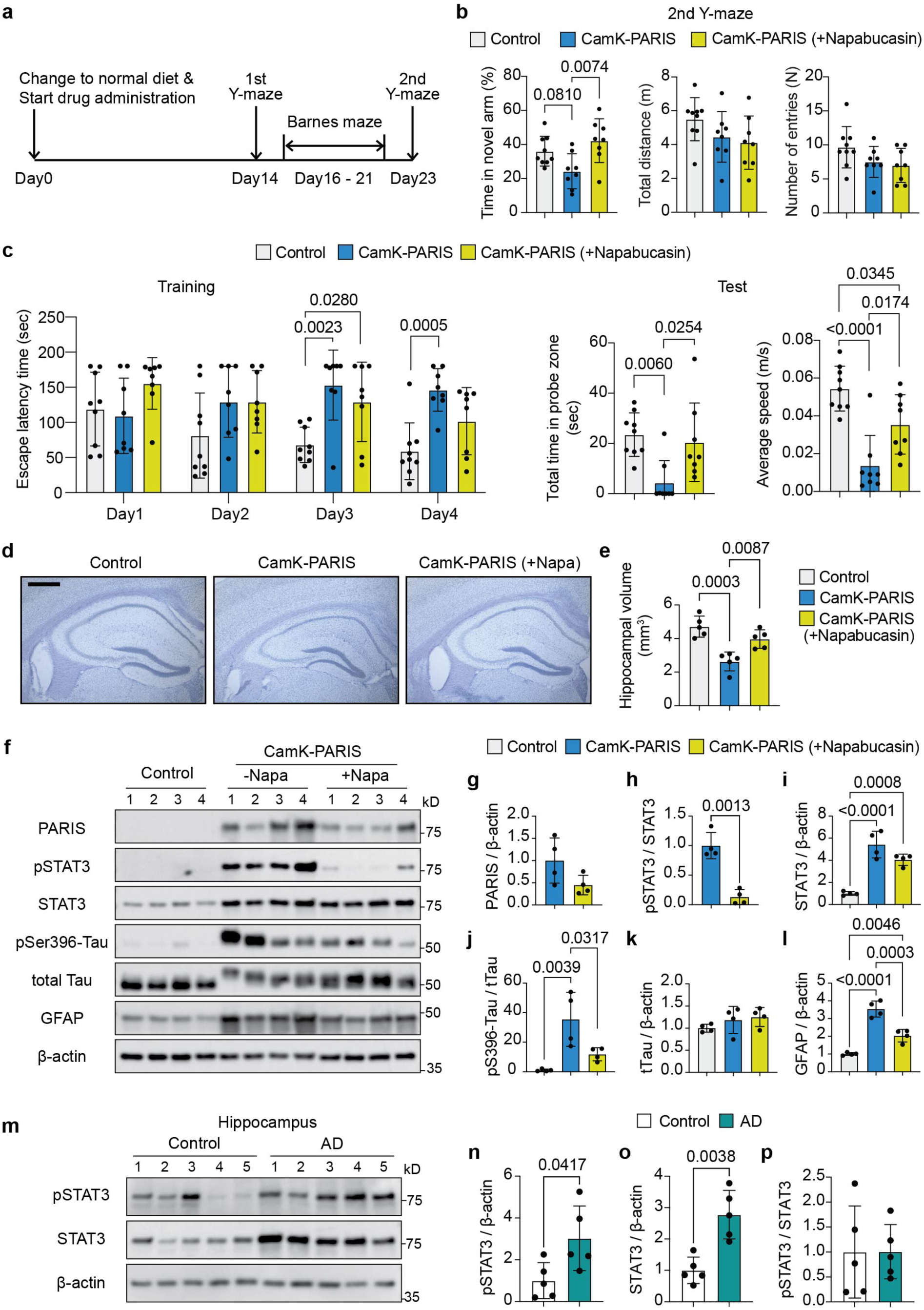
STAT3 inhibition reverses PARIS-driven behavioral deficits, tau phosphorylation, and hippocampal atrophy. **a**, Timeline for the Napabucasin administration and behavioral tests with control and CamK-PARIS mice. Normal diet and drug administration were started at 2 months of age. **b**, Percentage of time spent in the novel arm, total distance traveled, and arm entries in the second Y-maze test. Data are mean ± SEM (Control, *n* = 9; CamK-PARIS, *n* = 8; CamK-PARIS (+Napabucasin), *n* = 8). Statistical significance was assessed by one-way ANOVA followed by Tukey’s post hoc test. **c**, Quantification of escape latency during training, time spent in the target zone, and mean speed during the test session in the Barnes maze. Data are mean ± SEM (Control, *n* = 9; CamK-PARIS, *n* = 8; CamK-PARIS (+Napabucasin), *n* = 8). Statistical significance was assessed by one-way ANOVA followed by Tukey’s post hoc test. **d**, Representative Nissl-stained coronal sections of the hippocampus from mice. Scale bar, 500 µm. **e**, Quantification of hippocampal volume from images shown in **d**. Data are mean ± SEM (Control, *n* = 5; CamK-PARIS, *n* = 5; CamK-PARIS (+Napabucasin), *n* = 5). Statistical significance was assessed by one-way ANOVA followed by Tukey’s post hoc test. **f**, Immunoblot analysis of PARIS, pTyr705-STAT3, total STAT3, pSer396-tau, total tau, GFAP, and β-actin in hippocampal samples from CamK-PARIS mice. **g–l**, Quantification of pTyr705-STAT3, total STAT3, pSer396-tau, total tau, and GFAP levels from **f**. Data are mean ± SEM (Control, *n* = 4; CamK-PARIS, *n* = 4; CamK-PARIS (+Napabucasin), *n* = 4). Statistical significance was assessed by two-tailed *t*-test or one-way ANOVA followed by Tukey’s post hoc test. **m**, Immunoblot analysis of pTyr705-STAT3, total STAT3, and β-actin in hippocampal samples from control and AD patients. The same patient samples as in Fig. 1a were used. **n–p**, Quantification of pTyr705-STAT3 and total STAT3 levels from **m**. Data are mean ± SEM (control, *n* = 5; AD, *n* = 5). Statistical significance was assessed by two-tailed *t*-test. Comparisons with *P* ≤ 0.05 are marked on the graph.

In the Y-maze, no differences were detected between groups at 2 weeks after induction, but by 3 weeks CamK-PARIS mice exhibited impaired spatial memory, which was restored by napabucasin treatment (Fig. 6b and Extended Data Fig. 5a). This indicates that behavioral deficits emerge between 2–3 weeks of PARIS induction and are PARIS-dependent. We also conducted a Barnes maze to assess spatial learning and long-term memory. During training, group differences gradually increased, consistent with the emerging effects of PARIS overexpression (Fig. 6c). On the test day, CamK-PARIS mice treated with napabucasin spent significantly more time in the probe zone compared to untreated CamK-PARIS mice (Fig. 6c). A substantial difference in locomotor speed was also observed between groups (Fig. 6c), with analysis of average speed across training sessions showing a progressively widening disparity over time (Extended Data Fig. 5b). Because CamK-PARIS mice begin to exhibit premature mortality after approximately five weeks of PARIS induction, behavioral testing was constrained to a limited time window. Moreover, the daily oral gavage required for napabucasin administration imposed additional stress, making strongly aversive paradigms such as fear conditioning less suitable. We therefore employed the Barnes maze as a less stressful alternative, which also permits assessment of anxiety-like behaviors^45,46^. In this context, the differences in locomotor activity and exploratory behavior complicate the interpretation of escape latency as an isolated measure of spatial memory. Instead, these results more plausibly reflect heightened anxiety in CamK-PARIS mice, which was alleviated by STAT3 inhibition.

Given the behavioral rescue following STAT3 inhibition, we next assessed whether napabucasin treatment was associated with neuronal preservation by quantifying hippocampal volume. CamK-PARIS mice exhibited a marked reduction in hippocampal volume compared with control littermates, whereas napabucasin treatment attenuated this loss (Fig. 6d,e). In parallel, immunostaining for the presynaptic marker SYNAPSIN I and the postsynaptic marker PSD95 revealed a pronounced reduction in synaptic area in the hippocampus of CamK-PARIS mice, which was restored by napabucasin treatment (Extended Data Fig. 5c–e). Together, these findings indicate that STAT3 inhibition rescues neuronal loss–associated structural and synaptic alterations induced by PARIS overexpression.

At the molecular level, napabucasin treatment attenuated neuronal PARIS-induced signaling changes. In CamK-PARIS mice treated with napabucasin, phosphorylated STAT3 was reduced, while total STAT3 and PARIS levels remained unchanged (Fig. 6f–i). Napabucasin also restored the elevated levels of phosphorylated tau and GFAP (Fig. 6j–l). Consistent with these effects, immunostaining showed that napabucasin reversed the PARIS-induced increases in both GFAP and C3 intensities, while the C3-to-GFAP ratio remained unchanged (Extended Data Fig. 5f–i). This indicates that STAT3 inhibition rescues astrocytic expansion but does not further affect the activation state of individual astrocytes. Similarly, napabucasin rescued PARIS-induced microglial alterations, with reduced IBA1 intensity and, more importantly, a marked decrease in the proportion of CD16/CD32 positive cells within the IBA1 positive microglial population (Extended Data Fig. 5j–l). These findings indicate that STAT3 inhibition effectively attenuates PARIS-driven microglial activation.

Previous studies have reported increased STAT3 phosphorylation in the hippocampus of AD^47^. Consistent with these reports, we confirmed that both phosphorylated STAT3 and total STAT3 were elevated in the hippocampus of AD cases (Fig. 6m–o). In contrast, the ratio of phosphorylated STAT3 to total STAT3 remained unchanged (Fig. 6p), indicating that increased STAT3 signaling in AD is primarily driven by elevated STAT3 expression rather than a change in its phosphorylation status. Notably, the expression patterns of PARIS and total STAT3 closely paralleled each other (Fig. 1a, Fig. 6m), supporting the notion that PARIS may contribute to tau pathology in AD by driving STAT3 upregulation.

### Loss of *Paris* does not affect behavioral deficits and amyloid pathology in 5XFAD mice

To determine whether PARIS also modulates Aβ-driven pathology, we examined 5XFAD mice, which overexpress mutant human *APP* and *PSEN1* carrying five familial AD mutations^48^. In contrast to the PS19 tauopathy model, *Paris* deletion did not rescue cognitive impairments in 5XFAD mice. Y-maze, fear conditioning, and Morris water maze tests all revealed robust memory deficits in 5XFAD mice relative to wild-type littermates, and these impairments persisted in 5XFAD;*Paris*^⁻/⁻^ animals (Extended Data Fig. 6a–c). Consistent with these behavioral results, double labeling with Thioflavin S and Aβ antibody showed extensive plaque deposition in the hippocampus of 5XFAD mice, which was unaffected by *Paris* knockout (Extended Data Fig. 6d,e).

Immunoblotting likewise confirmed that elevated full-length APP and its C-terminal fragments (C99 and C83) were not altered by *Paris* deletion (Extended Data Fig. 6f–i). Notably, *Paris* knockout significantly reduced total STAT3, pTyr705-STAT3, and GFAP expression in 5XFAD mice (Extended Data Fig. 6f,j,k,m), paralleling the PARIS–STAT3–GFAP regulatory axis identified in PS19 mice. Despite this suppression of gliosis markers, there was no improvement in amyloid pathology or cognition, underscoring a mechanistic difference between tau- and amyloid-driven disease. As observed in PS19 mice, the ratio of pTyr705-STAT3 to total STAT3 remained unchanged, and IBA1 levels were unaffected (Extended Data Fig. 6l,n). Together, these findings indicate that while PARIS regulates STAT3 signaling and astrocytic reactivity across AD models, its pathological impact is specific to tauopathy rather than Aβ pathology.

## Discussion

Although tau-targeted therapeutic strategies are gaining increasing prominence in AD, the upstream mechanisms that initiate and amplify tau pathology remain incompletely defined^49^. In this study, we identify PARIS as a critical neuronal regulator of *STAT3* at the transcriptional level, establishing STAT3 as a direct effector of PARIS *in vivo*. Using complementary genetic and pharmacological approaches, we define a neuronal PARIS–STAT3 signaling pathway that links neuronal tau phosphorylation to glial activation and cognitive decline. These findings refine current views of AD pathogenesis by identifying an upstream neuronal determinant that connects tau pathology to reactive gliosis and provides mechanistic insight for therapeutic exploration.

STAT3 is recognized as a regulator of inflammatory responses^50,51^, and multiple STAT3 inhibitors are already undergoing clinical evaluation in oncology^52,53^. In the nervous system, STAT3 functions as a central mediator of neuroinflammation^54–56^, and its inhibition alleviates several AD-related pathologies, including tau hyperphosphorylation, amyloid deposition, synaptic dysfunction, and cognitive deficits^39,40,55,57^. Nevertheless, a previous study described an opposite effect, suggesting that STAT3 activation could exert protective actions in tauopathy^58^. In that work, the authors employed an acute viral model in which AAV-hTau was stereotaxically injected into the hippocampal CA3 region of 2-month-old wild-type mice, together with viral constructs for STAT3 downregulation or overexpression, and evaluated the mice one month later. While this design provides valuable insights into early signaling responses, the short duration may capture only partial aspects of tau-related pathology. In addition, total STAT3 levels in whole-brain extracts were reported to remain unchanged in that study, whereas in our 10-month-old PS19 mice, both phosphorylated and total STAT3 were markedly increased. Notably, even in the CamK-PARIS mice, in which neuronal PARIS expression was induced for three weeks, enhanced STAT3 expression and phosphorylation were accompanied by tau hyperphosphorylation and cognitive deficits. These findings suggest that model-dependent variables, including the duration and spatial extent of tau expression, may critically influence the observed role of STAT3. Moreover, maintaining appropriate STAT3 activity appears essential for neuronal integrity, as both excessive and insufficient activation can contribute to disease pathogenesis. A deeper understanding of how STAT3 signaling is regulated under distinct pathological contexts will be essential for defining its precise role in neurodegeneration.

Most studies on STAT3 activity in neurodegenerative models have focused on astrocytes, where dysregulation promotes astrogliosis and pro-inflammatory signaling^38,40,59,60^, leaving the contribution of neuronal STAT3 signaling to AD pathogenesis largely unexplored. Our results demonstrate that neuron-specific overexpression of PARIS significantly increased STAT3 expression and activation, thereby exacerbating tau phosphorylation and memory impairment. In contrast, astrocyte-specific PARIS overexpression did not induce tau pathology or behavioral deficits, indicating that PARIS promotes AD progression predominantly through neuronal STAT3 signaling. These observations highlight that neuronal perturbations alone can initiate downstream glial responses, placing our findings within the ongoing debate about the relative timing of neuronal and glial contributions to AD pathogenesis^61^.

AD has long been regarded as a process initiated by neuronal dysfunction and degeneration, which subsequently elicit glial responses^62^. This view is supported by observations that tau aggregation and synaptic loss occur within neurons and can secondarily trigger gliosis^22,23,63^. More recently, high-resolution approaches such as single-cell transcriptomics and spatial transcriptomics have suggested that glial alterations may emerge at presymptomatic stages, raising the possibility that astrocytes and microglia contribute to the initiation of pathogenesis rather than acting solely as downstream responders^19,64–67^. These findings have generated considerable debate, and reciprocal interactions between neurons and glia are now increasingly recognized as critical determinants of disease progression^68,69^. Within this context, our results support a model in which neuronal changes precede glial alterations within the PARIS–STAT3 pathway. Neuron-specific PARIS overexpression elevated STAT3, drove tau phosphorylation, and caused cognitive decline, followed by astrocytic and microglial activation. By contrast, astrocyte-specific PARIS overexpression increased STAT3 and GFAP but failed to induce tau pathology or behavioral impairments, underscoring that PARIS-driven tauopathy is neuron-dependent. Nonetheless, this neuron-first sequence is unlikely to be universally applicable, as other transcriptomic studies have also documented early glial alterations in specific stages and regions of AD^66,67,70^. Taken together, these results not only identify PARIS as a neuronal upstream driver of tauopathy but also highlight the PARIS–STAT3 axis as a mechanistic bridge linking neuronal dysfunction and reactive gliosis. As pharmacological strategies to modulate STAT3 signaling are already being explored in non-neurological diseases^52,53^, our findings provide a strong rationale for extending such approaches to AD and related tauopathies.

In addition to strategies targeting STAT3 directly, small molecules that inhibit PARIS activity have also shown neuroprotective potential. Farnesol, for instance, was reported to relieve PARIS-dependent repression of PGC1α, thereby mitigating mitochondrial dysfunction and neuronal loss *in vivo*^13^. Consistently, a subsequent study demonstrated that farnesol prevents aging-related muscle weakness in mice, further supporting the therapeutic potential of PARIS modulation^71^. Although *Paris* deletion in PS19 mice did not affect PGC1α expression, suggesting that this pathway may be less relevant in tauopathy, the study provides a proof-of-concept that PARIS function can be modulated to attenuate neurodegeneration. These findings raise the possibility that directly targeting PARIS to suppress its regulation of STAT3 may represent an effective therapeutic strategy for AD.

Our results also provide new insights into the functional versatility of PARIS as a transcription factor. PARIS belongs to the KRAB-ZNF family, characterized by an N-terminal KRAB domain and multiple C2H2 zinc finger motifs at the C-terminus^72,73^. Members of this family have been viewed as transcriptional repressors, primarily through recruitment of co-repressors that promote heterochromatin formation^72,73^.

However, accumulating evidence indicates that KRAB-ZNFs can also function as transcriptional activators depending on cellular context, interacting partners, and target genes^74,75^. Indeed, transcription factors are well known to switch between repressive and activating roles depending on the availability of co-repressors or co-activators^76,77^. Although PARIS was originally identified as a Parkin substrate and described as a repressor of *PGC1α* in Parkinson’s disease models, the same foundational study also reported that PARIS enhances *Pepck* promoter activity^15^, suggesting context-dependent activation. In the present study, we extend this concept by demonstrating that PARIS binds directly to the *STAT3* promoter and enhances its transcription in neurons, establishing *STAT3* as a novel target gene. These results highlight the functional versatility of PARIS and suggest that its activity as a transcriptional activator is mediated by yet unidentified cofactors. Elucidating which co-activators enable PARIS to drive *STAT3* transcription will be an important next step toward fully understanding the molecular mechanisms underlying this regulation.

Finally, the distinction between tau- and amyloid-driven mechanisms is an important consideration. Although *Paris* deletion reduced STAT3 and GFAP levels in 5XFAD mice, it did not ameliorate amyloid pathology or rescue cognitive deficits, indicating that PARIS plays a specific role in tau-mediated neurodegeneration. In line with this tau-specificity, the clinical observation of the APOE3 Christchurch carrier—who maintained cognitive resilience despite heavy amyloid deposition but minimal tau pathology—further supports the view that targeting tau-related mechanisms can improve cognitive outcomes in AD^9^. In this context, PARIS emerges as a promising therapeutic target for modifying disease progression.

In conclusion, our study identifies PARIS as a transcriptional activator of *STAT3* in neurons and establishes the PARIS–STAT3 axis as a mechanistic link between tau pathology and glial responses. By integrating behavioral, biochemical, and genetic evidence across multiple models, we show that PARIS amplifies tau pathology through neuronal STAT3 signaling and that STAT3 inhibition rescues these effects. Together, these findings refine our understanding of tau-driven disease initiation and position PARIS–STAT3 signaling as a promising target for therapeutic intervention in AD.

## Methods

### Animals

PS19 transgenic mice (P301S Tg mice, JAX: 008169) were obtained from Jackson Laboratory. *Paris* (*Znf746*) deficient mice were generated previously. PS19 mice were crossed with *Paris*^-/-^ mice to generate PS19;*Paris*^+/-^ mice, which were bred with *Paris*^+/-^ mice to obtain wild type, *Paris*^-/-^, PS19;*Paris*^+/+^, and PS19;*Paris*^-/-^ littermate mice. 5XFAD transgenic mice (JAX: 034840) were obtained from Jackson Laboratory and crossed with *Paris*^-/-^ mice using the same scheme as for the PS19 mice to obtain wild type, *Paris*^-/-^, 5XFAD;*Paris*^+/+^, and 5XFAD;*Paris*^-/-^ littermate mice. *CamKIIα*-*tTA* (JAX: 007004) and *GFAP-tTA* (JAX: 005964) mice were obtained from Jackson Laboratory were bred with *tetP*-*PARIS* mice, which were generated previously. All housing, breeding, and procedures were performed according to the NIH Guide for the Care and Use of Experimental Animals and approved by Johns Hopkins University Animal Care and Use Committee. Randomized mixed-gender cohorts were used for all animal experiments. There was no influence or association of gender with our findings.

### Human brain tissue

Fresh frozen hippocampus, entorhinal cortex, middle frontal gyrus, superior/middle temporal gyrus, and cerebellum tissues were obtained from the Division of Neuropathology, Department of Pathology at Johns Hopkins School of Medicine.

Information on CERAD score, Braak stage^78^, age at death, gender, race, and PMI of AD brain subjects is summarized in Supplementary Table 1.

### Tissue lysate preparation and western blot analysis

Tissues were homogenized the RIPA lysis buffer comprising 50mM Tris-HCl (pH 8.0), 150mM NaCl, 1% NP-40, 1% SDS, 0.5% sodium-deoxycholate and 1:100 Protease/Phosphatase Inhibitor Cocktail (Cell Signaling). Post homogenization, samples were kept at 4°C for 30 min. This homogenate was subsequently centrifuged at 16,000 g for 30 min, and the supernatant was collected for immunoblot. Protein levels were quantified and normalized with the Qubit Protein Assay Kit (Invitrogen). Laemmli Buffer plus β-mercaptoethanol was added to these samples. 20 mg of total protein lysates were separated using SDS polyacrylamide gels and transferred onto PVDF membranes (Bio-Rad). The blots were blocked with 5% nonfat dry milk in TBST (Cell Signaling) for 1 h and indicated primary antibodies were applied overnight at 4°C. Primary antibodies and working dilutions used were as follows: rabbit anti-PARIS antibody (Proteintech, 24543-1-AP, 1:2,000); mouse monoclonal anti-tau (TAU-5) antibody (Invitrogen, AHB0042, 1:4,000); rabbit monoclonal anti-tau antibody (Cell Signaling, 46687, 1:4,000); mouse monoclonal anti-phospho-tau (Ser202, Thr205) antibody (AT8) (Invitrogen, MN1020, 1:4,000); mouse monoclonal anti-phospho-tau (Thr181) antibody (AT270) (Invitrogen, MN1050, 1:4,000); rabbit polyclonal anti-phospho-Tau (Ser396) antibody (Invitrogen, 44-752G, 1:4,000); rabbit monoclonal anti-β-actin antibody (HRP Conjugate) (Cell Signaling, 5125, 1:5,000); rabbit polyclonal anti-FLAG antibody (Sigma-Aldrich, F7425, 1:4,000); mouse monoclonal anti-STAT3 antibody (Cell Signaling, 9139, 1:2,000); rabbit monoclonal phospho-STAT3 (Tyr705) antibody (Cell Signaling, Cat# 9145, 1:2,000); mouse monoclonal anti-glial fibrillary acidic protein (GFAP) antibody (Millipore Sigma, G3893,1:4,000); rabbit polyclonal anti-Iba1 antibody (FUJIFILM Wako Pure Chemical Corporation, 019-19741, 1:2,000); mouse monoclonal anti-APP antibody (Millipore Sigma, MAB348, 1:4,000); mouse monoclonal anti-APP C-Terminal Fragment Antibody (BioLegend, 802801, 1:2,000).

Target antigens were incubated with corresponding HRP-conjugated secondary antibodies (Cell signaling) and were visualized by ECL substrate (Thermo Scientific™).

ECL images were obtained with the Amersham Imager 600 (Cytiva) and analyzed with ImageJ software.

### CUT&Tag for DNA sequencing library

We followed the detailed protocol available at https://www.protocols.io/view/bench-top-cut-amp-tag-kqdg34qdpl25/v3?step=10. Briefly, to generate the Tn5-adapter complex, we annealed the Mosaic end adapter A and B oligonucleotides with their respective reverse complements, mixed the resulting products with pA-Tn5, and incubated the mixture for 1 hour. Hippocampal tissue from CamK-PARIS mice was homogenized, fixed with 0.1% formaldehyde, and cross-linking was halted with 1.25 M glycine. Concanavalin A-coated bead slurry was resuspended in Binding buffer, added to cells, and rotated. After clearing on a magnet stand, cells were resuspended in Antibody buffer, treated with primary antibody, and rotated overnight. Secondary antibody treatment followed, and bead washing was performed. The pA-Tn5 adapter complex was mixed, added to cells, and incubated. To stop tagmentation, DNA fragments were solubilized, purified, and PCR-amplified. The cycling program included denaturation and annealing steps. Post-PCR, SPRI bead purification and library assessment by capillary electrophoresis were conducted.

### CUT&Tag data analysis

Library quality was assessed using FastQC^79,80^. Paired-end reads were aligned to the reference genome using Bowtie2^81^ with the following parameters: --local --very-sensitive --no-mixed --no-discordant --phred33 -I 10 -X 700. Unmapped reads were removed with samtools^82^, and read pairs mapping to the same chromosome with a fragment length of less than 1,000 bp were retained. Peak calling was performed using SEACR^83^ with a normalized IgG control track and stringent threshold, and heatmaps were visualized with DeepTools^84^. Motif-based sequence analysis was carried out using the MEME Suite^85^, and peak annotation was performed with ChIPseeker^86^. Pathway analysis was conducted in WebGestalt^87^ using WikiPathways^88^, based on 1,139 genes corresponding to significant peaks within the top 5% of promoter-associated peaks. The frequency of unique genes represented in the top 10 significantly enriched pathways was calculated, and the most frequently involved genes were selected as candidate targets.

### Mouse tissue immunofluorescence analysis

Mice were perfused with PBS, and the brains were fixed with 4% paraformaldehyde at 4°C for 48 h followed by cryoprotection in 30% sucrose. 40 μm brain sections were blocked in 5% goat serum with 0.2% Triton X-100 in PBS for 1 h. Primary antibodies were prepared in 5% goat serum with 0.2% Triton X-100 in PBS incubated overnight at 4°C. Primary antibodies and working dilutions used were as follows: rabbit polyclonal anti-Synapsin I antibody (Millipore Sigma, AB1543, 1:1000); mouse monoclonal anti-PSD95 antibody (Novus Biologicals, NB300-556, 1:1000); mouse monoclonal anti-STAT3 antibody (Cell Signaling, 9139, 1:1,000); rabbit monoclonal phospho-STAT3 (Tyr705) antibody (Cell Signaling, Cat# 9145, 1:1,000); mouse monoclonal anti-glial fibrillary acidic protein (GFAP) antibody (Millipore Sigma, G3893, 1:2,000); rabbit polyclonal anti-Iba1 antibody (FUJIFILM Wako Pure Chemical Corporation, 019-19741, 1:1,000); rabbit polyclonal anti-complement C3 antibody (Invitrogen, PA5-21349, 1:1,000); rat monoclonal anti-CD16/CD32 antibody (Invitrogen, 14-0161-82, 1:1,000); mouse monoclonal anti-β-Amyloid antibody (6E10) (BioLegend, 803001, 1:1,000). After three-time washes in PBS which contains 0.2% Triton X-100 with the following day, Alexa-fluor conjugated secondary antibodies were used for 1 h at room temperature. Secondary antibodies and working dilutions used were as follows: Alexa Fluor 568 (Invitrogen, A-11031, 1:500); Alexa Fluor 488 (Invitrogen, A-11034, 1:500); Alexa Fluor 647 (Invitrogen, A-21247, 1:500). The sections were washed three times in PBS. ProLong Gold Antifade Mountant with DAPI (Invitrogen) was used to mount the sections. For ThioS staining, mouse tissue slices labeled with 6E10 antibody were placed on slides and allowed to dry overnight. The slides were then immersed in a 500 uM ThioS solution for 7 minutes followed by sequential washes in 100%, 90%, and 70% ethanol, and finally washed twice in PBS before mounting. Fluorescent images were acquired by confocal scanning microscopy using the Zeiss LSM 880 Microscope (Carl Zeiss). The selected area in the signal intensity range of the threshold was measured using ImageJ analysis.

### Immunohistochemistry and quantitative analysis

Brains were collected and fixed for 48 h in 4% paraformaldehyde and cryoprotected in 30% sucrose. Free-floating 40 μm sections were blocked with 5% horse serum with 0.2% Triton X-100 in PBS and incubated with an anti-phospho-tau (Ser202, Thr205) antibody (AT8) (Invitrogen, MN1020) in 1:1,000 dilution followed by incubation with biotin-conjugated anti-mouse antibody (Vectastain Elite ABC Kit). The sections were visualized with SigmaFast DAB peroxidase substrate (Sigma-Aldrich) and counterstained with Nissl stain (0.09% thionin). For image analysis, IHC Profiler was installed, and TIFF image files were opened using ImageJ. When the images were analyzed, and the percentage of the area occupied by the four ratings (High positive, Positive, Low Positive, Negative) was displayed. The AT8 positive area was determined by adding the percentage of high positive and positive.

### Hippocampal volumetric analysis

Sample brains were sectioned at 40 µm thickness, and one in every ten sections was selected for volumetric analysis, yielding a section sampling interval of 0.4 mm. Free-floating sections were mounted onto glass slides, air-dried, and subjected to Nissl staining using thionin (0.09%). Hippocampal areas were manually delineated on each section in ImageJ. Hippocampal volume was estimated by summing the measured areas from serial sections and multiplying by the section sampling interval (0.4 mm). Each data point represents the mean hippocampal volume per mouse calculated from all analyzed sections.

### Chromatin immunoprecipitation (ChIP) assay

ChIP was performed using a SimpleChIP^®^ Plus Enzymatic Chromatin IP Kit (Cell Signaling Technology). Briefly, hippocampus from CamK-PARIS mice were crosslinked with 1.5% formaldehyde in PBS which contains Protease Inhibitor Cocktail (PIC) at RT for 20 min and stop cross-linking by adding 2.5 M glycine at RT for 5 min and washed with PBS two times. Tissues were disaggregated into single-cell suspension using a handy homogenizer (VWR, Radnor, PA, USA). Chromatin was incubated with micrococcal nuclease at 37°C for 20 min with frequent mixing by inversion to digest DNA which was followed by sonication to break nuclear membrane. The supernatants were immunoprecipitated by being incubated with overnight at 4°C with 1 μg of anti-PARIS or anti-IgG. The next day, 30 µl of Protein G Magnetic Beads were added to each IP reaction and incubate for 2 hours at 4°C with rotation and then were washed using low salt wash buffer and high salt wash buffer. Chromatin was eluted by ChIP elution buffer for 30 min at 65 °C with gentle vortex mixing and crosslinks were reversed by treatment with 5 M NaCl and proteinase K at 65 °C for 2 h. ChIP DNA was purified and quantified by quantitative real-time PCR. IP efficiency was calculated manually using the equation (Percent Input = 2% x 2^(C[T]^ ^2%Input^ ^Sample-C[T]^ ^IP^ ^Sample)^). Primer sequences for PCR are listed in Supplementary Table 2.

### Real-time PCR

Total RNA was isolated from hippocampus using the RNeasy Plus Mini Kit (Qiagen) according to the manufacturer’s instructions. RNA concentrations were measured with a NanoDrop 2000 spectrophotomer (Biotek, Winooski, VT, USA). Total RNA (500 ng) was reverse-transcribed to cDNA with the GoScript Reverse Transcription Mix (Promega, Madison, WI, USA) for quantitative RT-PCR (qPCR) with a Viia7 Real-Time PCR System (Applied Biosystems, Waltham, MA, USA) using SYBR Green Master Mix (Applied Biosystems). Relative gene expression was quantified after normalizing to β-actin expression using the relative standard curve method. Primer sequences used are listed in Supplementary Table 3.

### Behavioral Tests

#### Novel object recognition test (NORT)

The NORT was performed according to the method described previously^30^. NORT were carried out in a box (30 cm width x 30 cm depth x 30 cm height) with polyvinyl plastic which was covered by black paper. Prior to conducting the NORT, all mice underwent a habituation stage in which they were individually introduced to the test box for a 5-minute period without any objects present. Subsequently, mice were placed into the test box containing two identical wooden blocks. They were allowed to freely explore these objects for a duration of 10 minutes. The time spent by each mouse exploring each object was recorded during this training session. Twenty-four hours following the training session, mice were reintroduced to the test box. One of the familiar objects from the training session was replaced with a novel object. The mice were given another 10-minute session to explore the objects. The time spent by each mouse exploring both the novel and familiar objects was recorded during this test session. The behavior of exploration was defined as when a mouse was observed sniffing an object. The objects and test box were cleaned with 10% ethanol after each session. Data are expressed in percentage terms of novel object recognition time (time percentage = total time spent with novel object/[total time spent with novel object + total time spent with familiar object] x 100).

#### Y-maze test (spatial recognition)

The maze with three equal angles between all arms, which were 40 cm long and 10 cm wide with 15 cm high walls was used. The maze was constructed using opaque polyvinyl plastic. The training phase began by placing a mouse at the entrance of one of the two open arms, allowing it to explore the maze freely for 5 minutes. During this phase, one of the three arms remained blocked. Once the training phase finished, the mouse was returned to its home cage. After 45 minutes interval, the obstruction in one of the maze arms was removed, allowing access to all three arms. The mouse was then placed at the entrance of one of the familiar arms at the beginning of the test phase. During this test phase, the mouse was given 5 minutes to freely explore the maze. The percentage of time spent by the mouse in the novel arm during the initial 2 minutes was measured. Thorough maze cleaning followed each session to eliminate olfactory cues.

#### Fear conditioning test

Fear conditioning test was conducted with a white light on in a conditioning chamber (Med Associates Inc) with transparent walls and a floor made of stainless-steel rods. One day before shock presenting, allow the mouse to acclimate to the experimental chamber for 10 minutes without any stimuli or shocks as a habituation phase. In shock presentation phase, mouse is placed in the chamber and present with a tone for 2 seconds then deliver an electric shock (0.5 mA) for 2 seconds. A total of four tone-shock pairings (120 s, 340 s, 580 s, 780 s) are applied for 1010 seconds. The chamber was cleaned with 10% ethanol after each session. One day later, during the cue test phase, mice were placed to the conditioning chamber and tones were applied without electric shock at the same intervals as in the shock presentation phase. For each tone, the total freezing time up to that point was measured. To exclude visual and olfactory effects, the white light in the chamber was turned off, the sides and floor were covered with acrylic plates, and the chamber was cleaned with 0.25% acetic acid every session.

#### Morris water maze test

The Morris water maze test^89^ was conducted using a circular pool measuring 130 cm in diameter and 50 cm in height, containing four distinct inner cues on the surface. The pool was filled with water and a nontoxic, water-soluble white dye. A submerged platform, placed 1 cm below the water’s surface, remained invisible at the water level. The pool was partitioned into four equal quadrants. Within one of these quadrants, a black platform measuring 9 cm in diameter and 15 cm in height was centrally located. Mouse movements, from their starting positions to the platform, were tracked and recorded using a video tracking system (ANY-Maze). On the day preceding testing, mice were subjected to swim training for 60 s in the absence of the platform. For five consecutive days, mice were subjected to three trial sessions per day, with an inter-trial interval of 1 hour. The escape latency, denoting the time taken to locate the platform, was recorded. This parameter was averaged for each session of trials and for each individual mouse. Once a mouse located the platform, it was allowed to remain on it for 10 s. If the mouse was unable to locate the platform within 60 s, it was placed on the platform for 10 s and then returned to its cage. On day 7 of testing, a probe trial was conducted. During this trial, the platform was removed from the pool, and the mice were given 60 s to explore the pool in the absence of the platform.

#### Barnes maze test

The Barnes maze test was performed according to the method described previously^45^. Briefly, the Barnes maze test comprises three main phases: habituation, training, and probe trial. During habituation phases, mouse was introduced to the maze with an escape tunnel, exploring until entering the tunnel or for a maximum of 5 minutes. The maze and tunnel were cleaned afterward. In the acquisition training phase, two daily trials last a maximum of 3 minutes each, with the tunnel’s position fixed relative to room cues and the starting location varied among quadrants. Training lasts for 4 days, and the escape latency of each mouse was measured. The next day, the probe trial phase lasted for 3 minutes, and the time spent in the quadrant where the tunnel was located was measured.

### Statistical analysis

All statistical analyses were performed using GraphPad Prism 10. Data are presented as mean ± SEM, and the number of biologically independent animals (*n*) is indicated in each figure legend. For histological and immunostaining experiments, multiple sections were analyzed per mouse, and the average value per animal was used for statistical comparisons. Animals were randomly assigned to experimental groups, and behavioral testing and image quantification were performed under blinded conditions. For comparisons between two groups, unpaired two-tailed Welch’s *t*-tests were used, and for comparisons among more than two groups, one-way ANOVA followed by Tukey’s post hoc test was applied. No statistical methods were used to predetermine sample size, but group sizes were similar to those generally employed in the field. All key findings were confirmed in at least two independent cohorts of animals, and reproducibility of molecular, histological, and behavioral experiments was validated across multiple litters and independent experiments.

## Data availability

All data supporting the findings of this study are available within the paper and Extended Data. Raw/source data have been deposited in the Dryad data repository and are publicly accessible (https://doi.org/10.5061/dryad.70rxwdcbz). The CUT&Tag chromatin profiling data have been deposited in the Gene Expression Omnibus (GEO) under accession number GSE309968. There are no restrictions on data availability. Source data are provided with this paper.

## Acknowledgements

We thank the members of the Dawson Laboratory for helpful discussions and technical assistance. This work was supported in part by funding administered through the Dawson Lab and it was supported in part by the Freedom Together Foundation, the Thome Memorial Foundation and the Alzheimer’s Association Zenith Fellows Award. T.M.D. is the Leonard and Madlyn Abramson Professor in Neurodegenerative Diseases. Human brain postmortem tissues availability was supported by the JHU Alzheimer’s Disease Research Center (NIH P30AG0066507) and the BIOCARD Study (U19AG033655).

## Author contributions

T.M.D. and V.L.D conceived the study. J.Y.S. designed and performed the experiments with assistance from F.A., H.J. and S.H. J.P. analyzed the CUT&Tag sequencing results. J.R.O. and J.C.T. provided the postmortem human brain tissue. J.Y.S. wrote the manuscript with editing by T.M.D., V.L.D. and S.U.K. T.M.D., V.L.D. and S.U.K. supervised the project.

---

**Extended Data Fig. 1.**
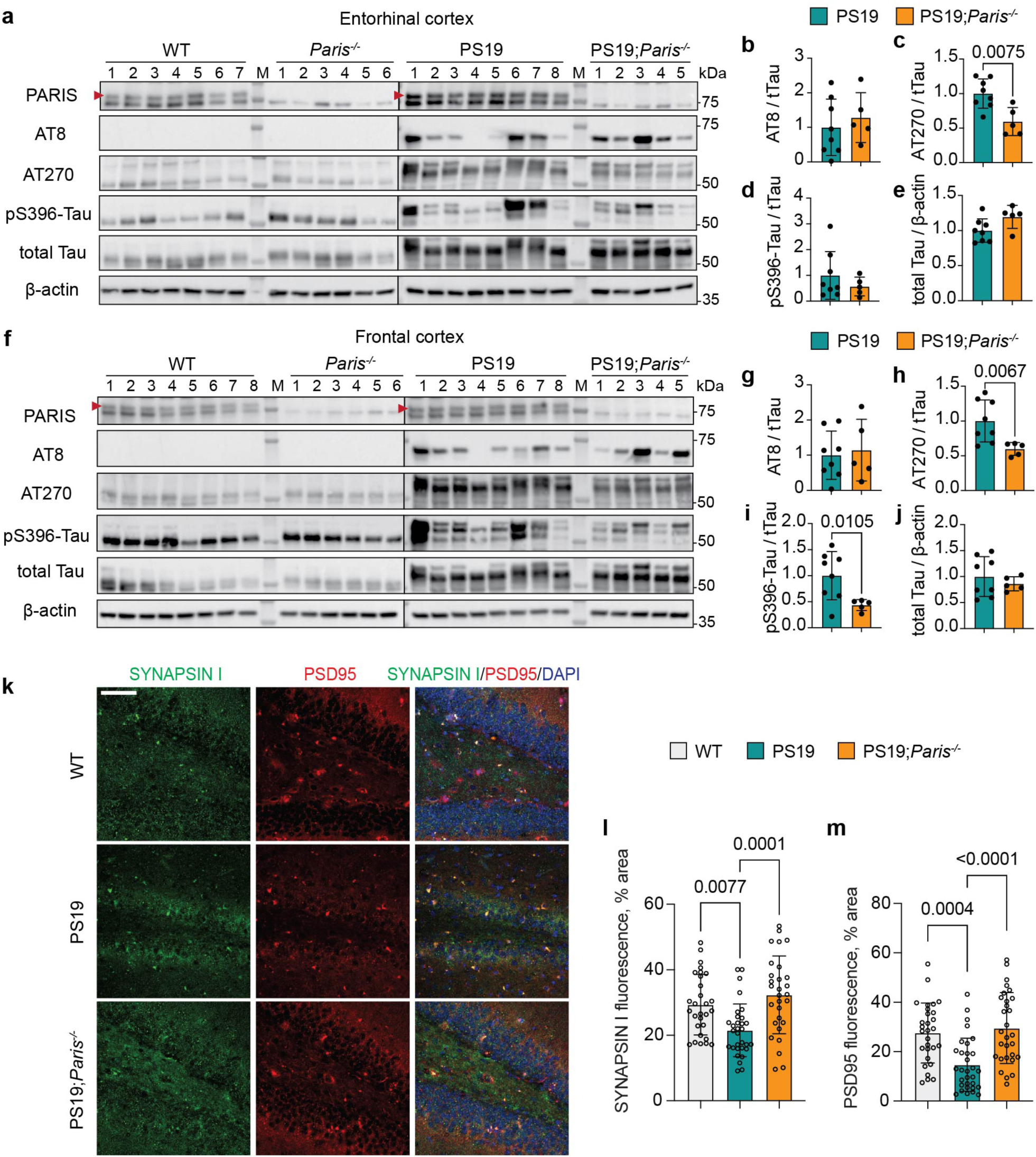
Loss of *Paris* reduces cortical tau phosphorylation and restores synaptic markers in PS19 mice. **a**, Immunoblot analysis of PARIS, pSer202/pThr205-tau (AT8), pThr181-tau (AT270), pSer396-tau, total tau, and β-actin in entorhinal cortex samples from 10-month-old mice. The location of PARIS is indicated by arrows. **b–e**, Quantification of total tau, AT8, AT270, and pSer396-tau levels from **a**. Data are mean ± SEM (PS19, *n* = 8; PS19;*Paris*^-/-^, *n* = 5). Statistical significance was assessed by two-tailed *t*-test. **f**, Immunoblot analysis of PARIS, pSer202/pThr205-tau (AT8), pThr181-tau (AT270), pSer396-tau, total tau, and β-actin in frontal cortex samples from 10-month-old mice. The location of PARIS is indicated by arrows. **g–j**, Quantification of total tau, AT8, AT270, and pSer396-tau levels from **f**. Data are mean ± SEM (PS19, *n* = 8; PS19;*Paris*^-/-^, *n* = 5). Statistical significance was assessed by two-tailed *t*-test. **k**, Representative immunofluorescence images of SYNAPSIN I (green), PSD95 (red), and DAPI (blue) in the hippocampus of 10-month-old mice. Scale bar, 50 µm. **l**,**m**, Quantification of SYNAPSIN I–positive area and PSD95–positive area from **k**. Data are mean ± SEM (WT, *n* = 29; PS19, *n* = 31; PS19;*Paris*^-/-^, *n* = 29; images from 5–8 sections per mouse and 5 mice per group). Statistical significance was assessed by one-way ANOVA followed by Tukey’s post hoc test. Comparisons with *P* ≤ 0.05 are marked on the graph.

**Extended Data Fig. 2.**
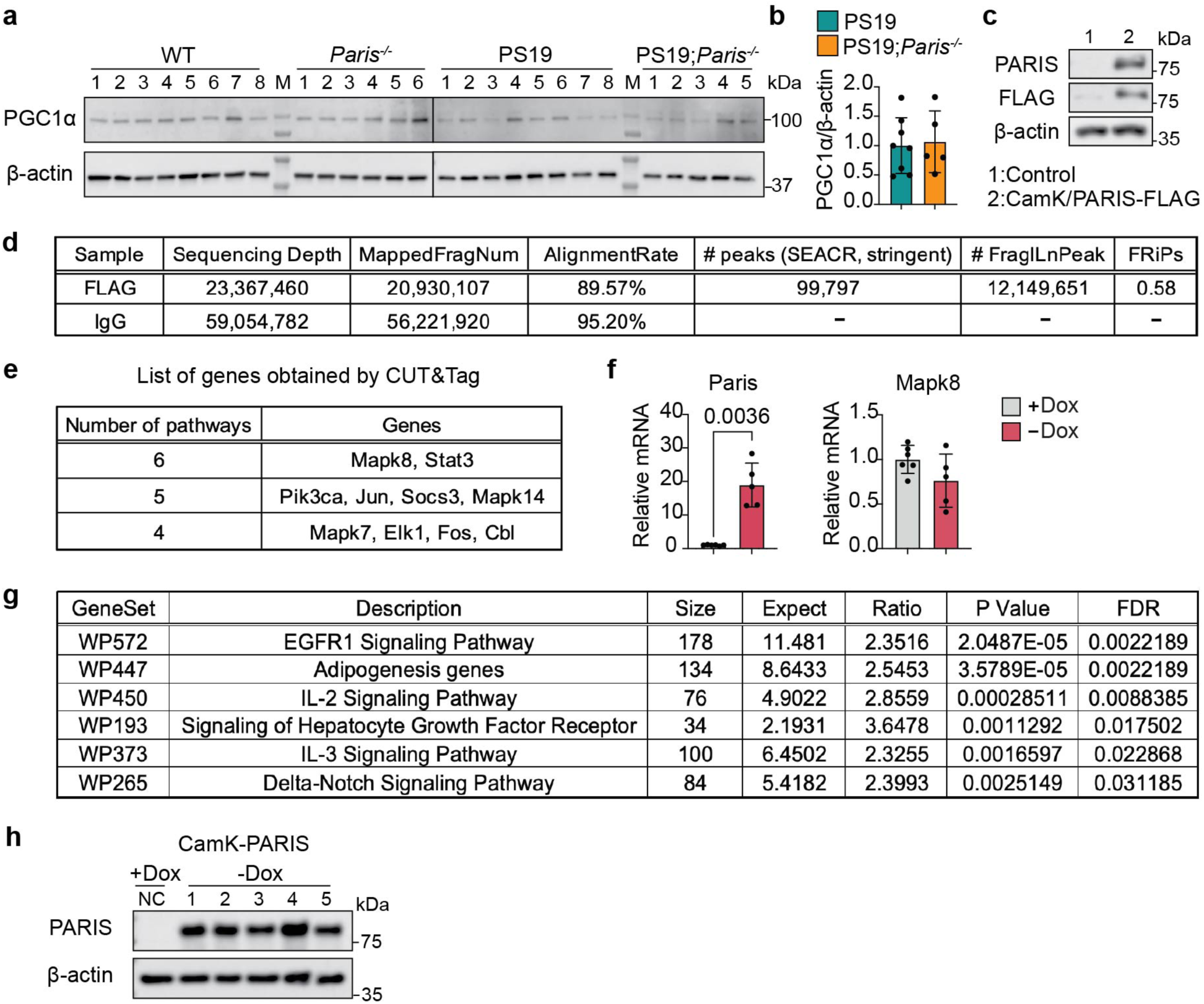
The results of CUT&Tag analysis and target gene validation. **a**, Immunoblot analysis of PGC-1α and β-actin in hippocampal samples from mice. The same mouse samples as in Fig. 2d were used. **b**, Quantification of PGC-1α from a. Data are mean ± SEM (PS19, *n* = 8; PS19;*Paris*^-/-^, *n* = 5). No significant differences were detected between groups (two-tailed *t*-test). **c**, Immunoblot analysis of FLAG-tagged PARIS and β-actin in hippocampal samples from the CamK-PARIS mouse used for CUT&Tag. **d**, The results of sequencing using CUT&Tag. Fragment proportion in peak regions (FRiPs) = (# FragInPeak / MappedFragNum). **e**, List of genes involved in high frequency in important pathways obtained by CUT&Tag. **f**, Relative mRNA expression *of Paris and Mapk8* normalized to β-actin, measured by quantitative RT-PCR. Hippocampal tissues from CamK-PARIS mice were used. Data are mean ± SEM (+Dox, *n* = 6; −Dox, *n* = 5). Statistical significance was assessed by two-tailed *t*-test. **g**, STAT3-related pathways obtained from CUT&Tag analysis. **h**, Immunoblot to confirm the expression of PARIS in the hippocampus of CamK-PARIS mice used for ChIP assay. Comparisons with *P* ≤ 0.05 are marked on the graph.

**Extended Data Fig. 3.**
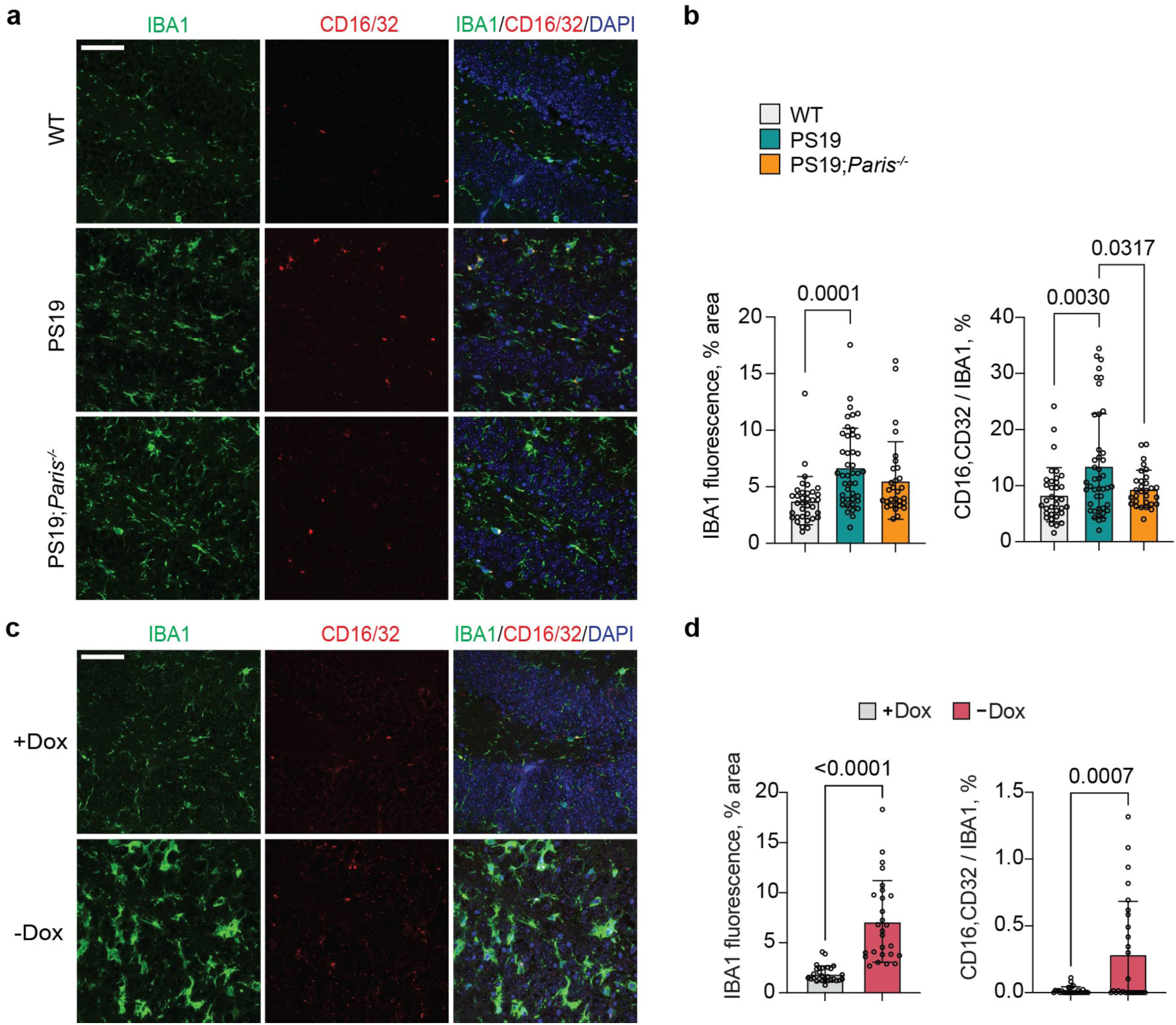
PARIS increases activation of microglia. **a**, Representative co-immunostaining images of IBA1 (green), CD16/CD32 (red), and DAPI (blue) in the hippocampus of 10-month-old PS19 mice. Scale bar, 50 µm. **b**, Quantification of IBA1 and relative CD16/CD32 fluorescence from **a**. Data are mean ± SEM (WT, *n* = 35; PS19, *n* = 43; PS19;*Paris*^-/-^, *n* = 30; images from 5–8 sections per mouse and 5–6 mice per group). Group differences were assessed by one-way ANOVA followed by Tukey’s post hoc test. **c**, Representative co-immunostaining images of IBA1 (green), CD16/CD32 (red), and DAPI (blue) in the hippocampus of CamK-PARIS mice. Scale bar, 50 µm. **d**, Quantification of IBA1 and relative CD16/CD32 fluorescence from **c**. Data are mean ± SEM (+Dox, *n* = 28; −Dox, *n* = 27; images from 4–6 sections per mouse and 5 mice per group). Statistical significance was assessed by two-tailed *t*-test. Comparisons with *P* ≤ 0.05 are marked on the graph.

**Extended Data Fig. 4.**
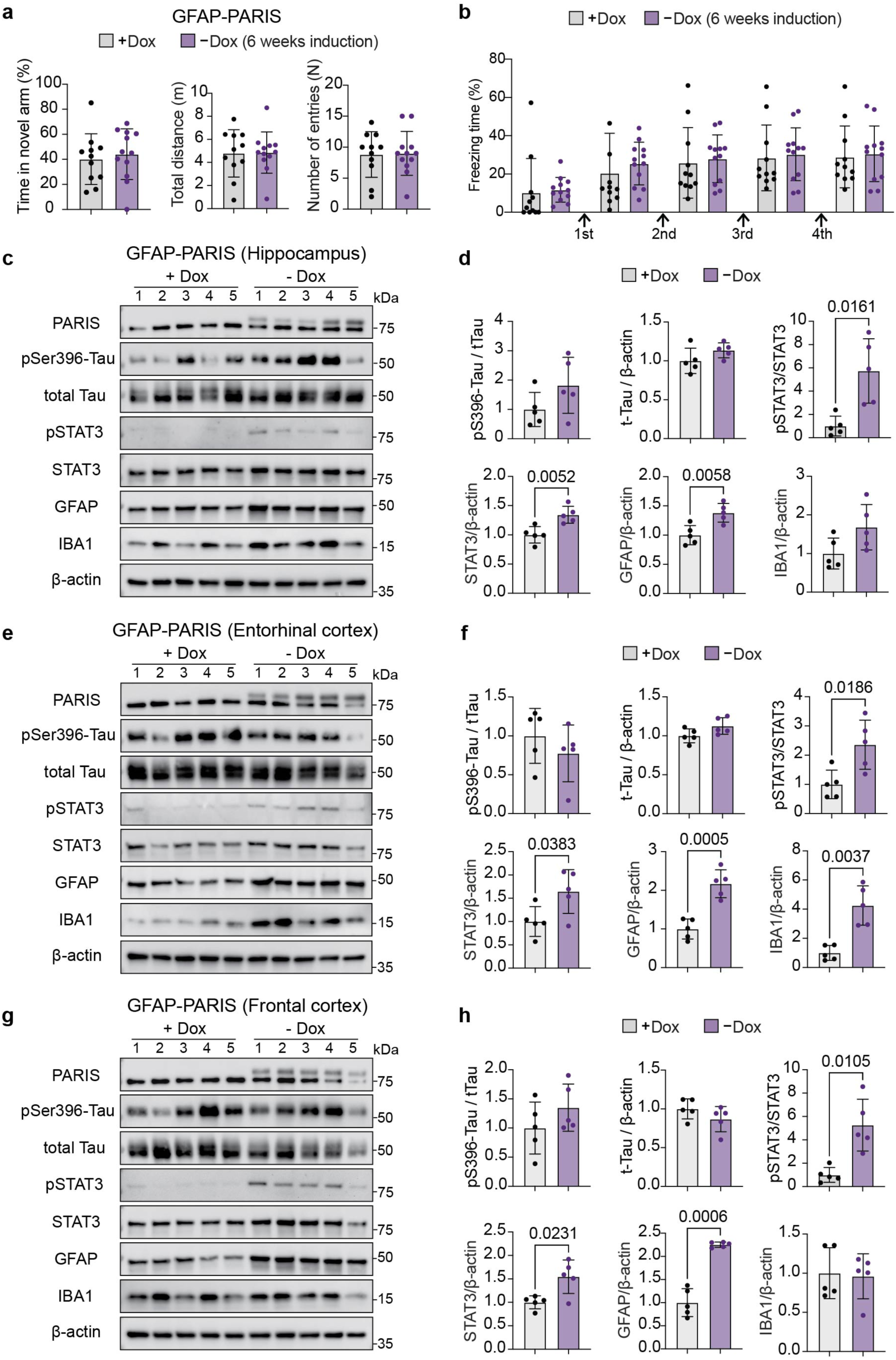
Astrocyte-specific PARIS overexpression does not affect mouse behavior and tau phosphorylation. PARIS expression was induced for 6 weeks by doxycycline withdrawal at 2 months of age. **a**, Percentage of time spent in the novel arm, total distance, and number of arm entries in the Y-maze test. Data are mean ± SEM (+Dox, *n* = 11; −Dox, *n* = 12). Statistical significance was assessed by two-tailed *t*-test. **b**, Percentage of freezing time at baseline and after four stimulus tones in the fear conditioning test with GFAP-PARIS mice. Data are mean ± SEM (+Dox, *n* = 11; −Dox, *n* = 12). Statistical significance was assessed by two-tailed *t*-test. **c**, Immunoblot of PARIS, pSer396-tau, total tau, pTyr705-STAT3, STAT3, GFAP, IBA1, and β-actin in the hippocampus of GFAP-PARIS mice. PARIS expression was induced for 6 weeks by doxycycline withdrawal at 2 months of age. **d**, Quantification of pSer396-tau, total tau, pTyr705-STAT3, STAT3, GFAP, and IBA1 from **c**. Data are mean ± SEM (+Dox, *n* = 5; −Dox, *n* = 5). Statistical significance was assessed by two-tailed *t*-test. **e**, Immunoblot of PARIS, pSer396-tau, total tau, pTyr705-STAT3, STAT3, GFAP, IBA1, and β-actin in entorhinal cortex samples from GFAP-PARIS mice. **f**, Quantification of pSer396-tau, total tau, pTyr705-STAT3, STAT3, GFAP, and IBA1 from **e**. Data are mean ± SEM (+Dox, *n* = 5; −Dox, *n* = 5). Statistical significance was assessed by two-tailed *t*-test. **g**, Immunoblot of PARIS, pSer396-tau, total tau, pTyr705-STAT3, STAT3, GFAP, IBA1, and β-actin in frontal cortex samples from GFAP-PARIS mice. **h**, Quantification of pSer396-tau, total tau, pTyr705-STAT3, STAT3, GFAP, and IBA1 from **g**. Data are mean ± SEM (+Dox, *n* = 5; −Dox, *n* = 5). Statistical significance was assessed by two-tailed *t*-test. Comparisons with *P* ≤ 0.05 are marked on the graph.

**Extended Data Fig. 5.**
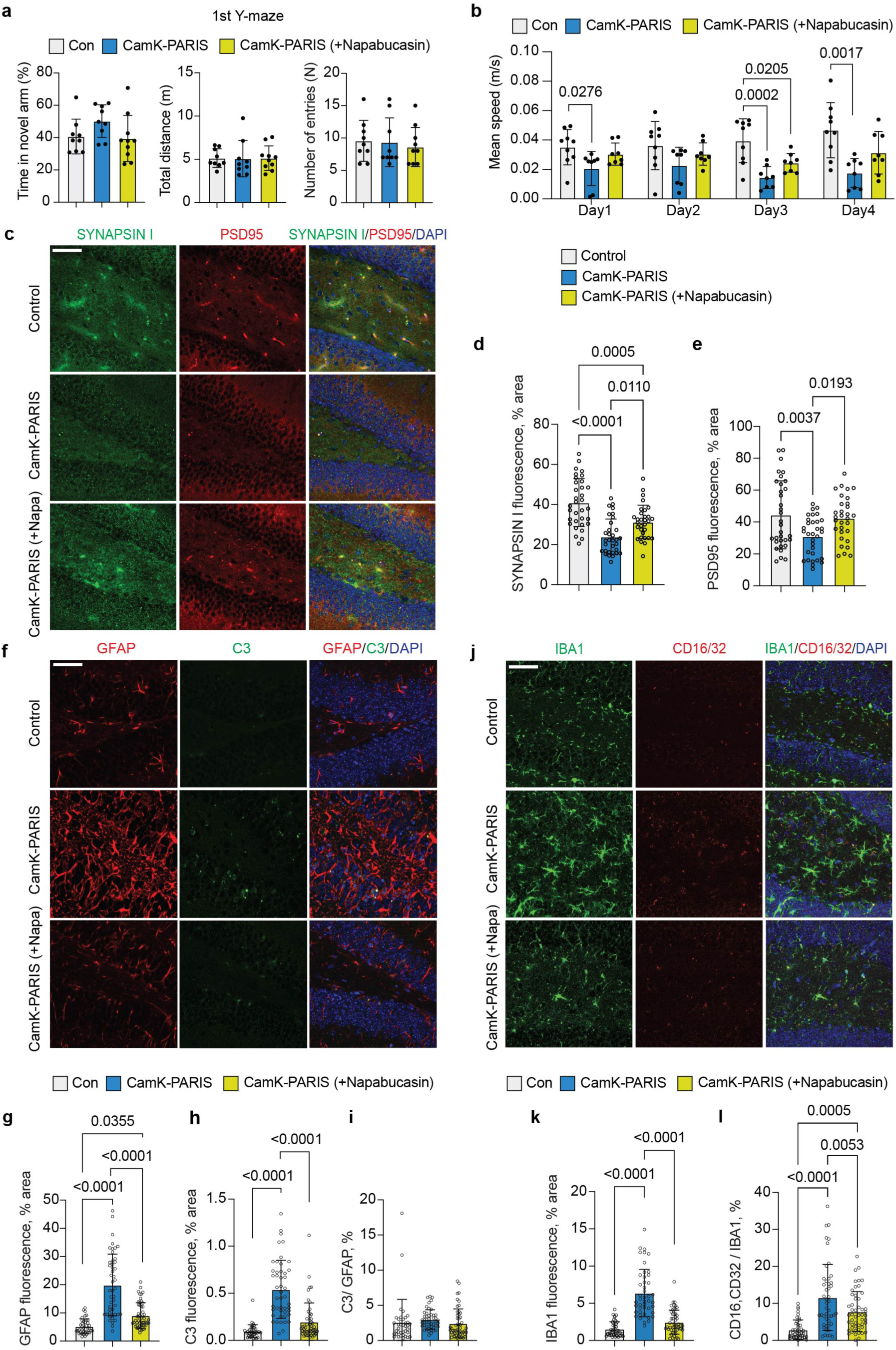
STAT3 inhibition restores synaptic markers and suppresses glial activation in CamK-PARIS mice. **a**, Percentage of time in the novel arm, total distance traveled, and arm entries in the first Y-maze test. Data are mean ± SEM (Control, *n* = 9; CamK-PARIS, *n* = 9; CamK-PARIS (+Napabucasin), *n* = 10). Group differences were assessed by one-way ANOVA followed by Tukey’s post hoc test. **b**, Mean speed during Barnes maze training. Data are mean ± SEM (Control, *n* = 9; CamK-PARIS, *n* = 8; CamK-PARIS (+Napabucasin), *n* = 8). Group differences were assessed by one-way ANOVA followed by Tukey’s post hoc test. **c**, Representative immunofluorescence images of SYNAPSIN I (green), PSD95 (red), and DAPI (blue) in the hippocampus of mice. Scale bar, 50 µm. **d**,**e**, Quantification of SYNAPSIN I–positive area and PSD95–positive area from **c**. Data are mean ± SEM (Control, *n* = 33; CamK-PARIS, *n* = 32; CamK-PARIS (+Napabucasin), *n* = 31; images from 5–8 sections per mouse and 5 mice per group). Statistical significance was assessed by one-way ANOVA followed by Tukey’s post hoc test. Comparisons with *P* ≤ 0.05 are marked on the graph. **f**, Representative co-immunostaining images of GFAP (red), C3 (green), and DAPI (blue) in the hippocampus of control and CamK-PARIS mice. Scale bar, 50 µm. **g–i**, Quantification of GFAP and relative C3 fluorescence from **f**. Data are mean ± SEM (Control, *n* = 38; CamK-PARIS, *n* = 47; CamK-PARIS (+Napabucasin), *n* = 51; images from 7–12 sections per mouse and 5 mice per group). Statistical significance was assessed by one-way ANOVA followed by Tukey’s post hoc test. **j**, Representative co-immunostaining images of IBA1 (green), CD16/CD32 (red), and DAPI (blue) in the hippocampus of control and CamK-PARIS mice. Scale bar, 50 µm. **k**,**l**, Quantification of IBA1 and relative CD16/CD32 fluorescence from **j**. Data are mean ± SEM (Control, *n* = 43; CamK-PARIS, *n* = 45; CamK-PARIS (+Napabucasin), *n* = 50; images from 7–10 sections per mouse and 5 mice per group). Statistical significance was assessed by one-way ANOVA followed by Tukey’s post hoc test. Comparisons with *P* ≤ 0.05 are marked on the graph.

**Extended Data Fig. 6.**
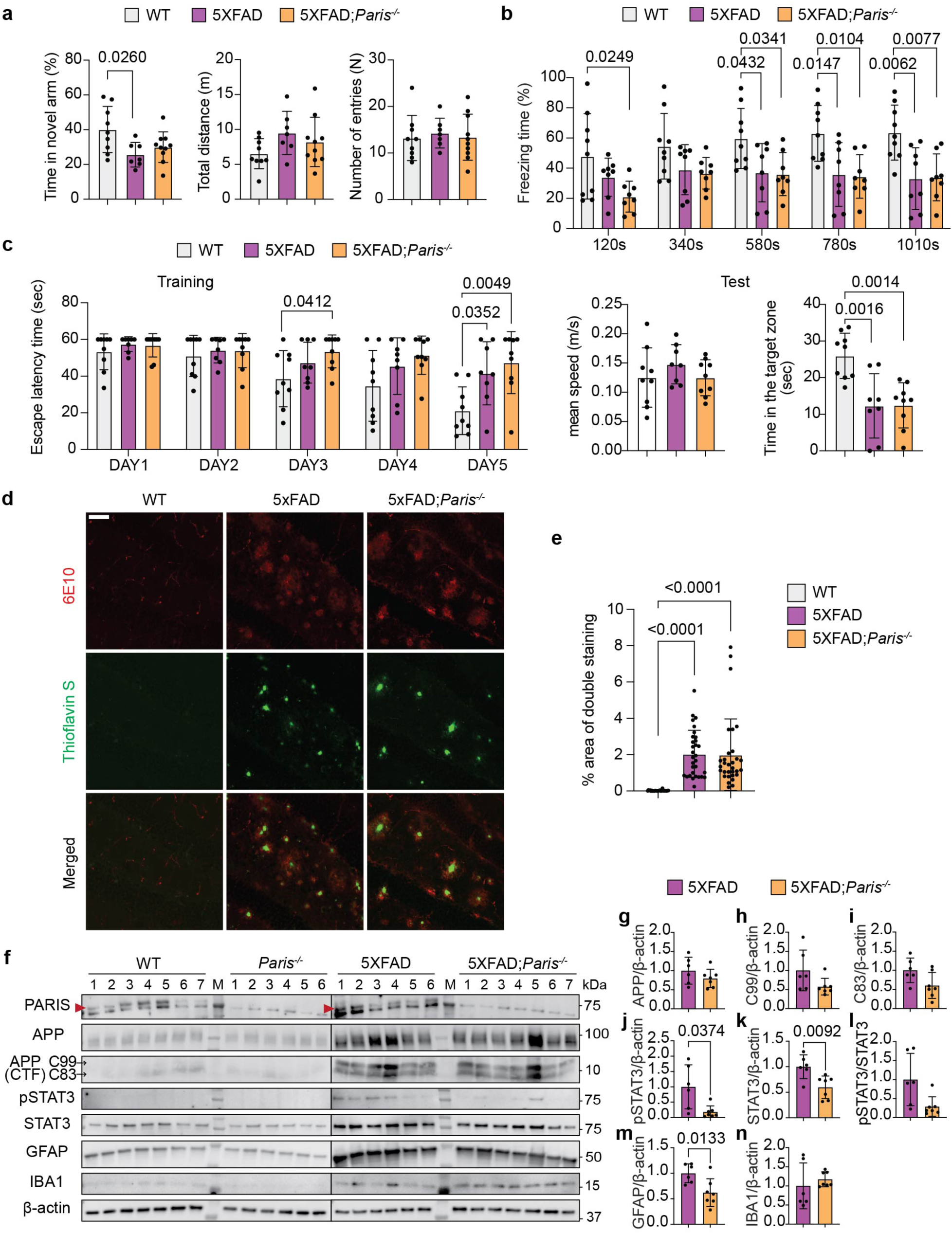
*Paris* knockout does not rescue behavioral deficits and pathological markers in 10-month-old 5XFAD mice. **a,** Percentage of time spent in the novel arm, total distance and number of arm entries in the Y-maze test (one-way ANOVA followed by Tukey’s post hoc test; WT, *n* = 9; 5XFAD, *n* = 7; 5XFAD;*Paris*^-/-^, *n* = 10). **b**, Percentage of freezing time at baseline and after four stimulus tones in 5XFAD mice in the fear conditioning test (one-way ANOVA followed by Tukey’s post hoc test; WT, *n* = 9; 5XFAD, *n* = 8; 5XFAD;*Paris*^-/-^, *n* = 8). **c**, Quantification of time to find the hidden platform during the training session and time spent in the target zone during the test day with the hidden platform removed in Morris water maze (one-way ANOVA followed by Tukey’s post hoc test; WT, *n* = 9; 5XFAD, *n* = 8; 5XFAD;*Paris*^-/-^, *n* = 9). **d**, Representative double-staining images of Aβ antibody (red) and Thioflavin-S (green) in the hippocampus of 10-month-old mice. Scale bar, 50 µm. **e**, Quantification of double-stained area with Aβ antibody (6E10) and Thioflavin-S from **d**. Data are mean ± SEM (WT, *n* = 26; 5XFAD, *n* = 31; 5XFAD;*Paris*^-/-^, *n* = 31; images from 4–6 sections per mouse and 6 mice per group). Statistical significance was assessed by one-way ANOVA followed by Tukey’s post hoc test. **f**, Immunoblot analysis of PARIS, APP, C99, C83, pTyr705-STAT3, STAT3, GFAP, IBA1, and β-actin in hippocampal samples from 10-month-old mice. The location of PARIS is indicated by arrows. **g–n**, Quantification of APP, C99, C83, pTyr705-STAT3, STAT3, GFAP, and IBA1 from **f**. Data are mean ± SEM (5XFAD, *n* = 6; 5XFAD;*Paris*^-/-^, *n* = 7). Statistical significance was assessed by two-tailed *t*-test. Comparisons with *P* ≤ 0.05 are marked on the graph.

**Supplementary Table 1.** Case information of control and AD patients brains (related to Fig. 1). Available information on CERAD score, Braak stage, age at death, sex (M = male, F = female), race (W = White, B = Black), postmortem delay (PMI). Hippo = hippocampus, ERC = entorhinal cortex, MFG = middle frontal gyrus, SMTG = superior/middle temporal gyrus, CB = cerebellum.

|  | FDX | CERAD | Braak | Age | Sex | Race | PMD | Hippo | ERC | MFG | SMTG | CB |
| --- | --- | --- | --- | --- | --- | --- | --- | --- | --- | --- | --- | --- |
| 1 | AD | C | 6 | 80 | F | W | 7 |  | ✓ |  |  |  |
| 2 | AD | C | 6 | 83 | F | W | 15.5 |  |  | ✓ |  |  |
| 3 | AD | C | 6 | 85 | F | W | 18 | ✓ |  |  |  |  |
| 4 | AD | C | 5 | 88 | F | W | 8 |  |  |  |  | ✓ |
| 5 | AD | C | 6 | 84 | F | W | 20 |  |  | ✓ |  | ✓ |
| 6 | AD | C | 6 | 83 | F | W | 19 |  |  | ✓ |  | ✓ |
| 7 | AD | C | 6 | 83 | F | W | 11 |  |  |  |  | ✓ |
| 8 | AD | C | 6 | 88 | M | W | 4 |  | ✓ |  |  |  |
| 9 | AD | C | 6 | 85 | F | W | 18 | ✓ | ✓ |  |  |  |
| 10 | AD | C | 6 | 86 | F | W | 14 | ✓ |  |  |  |  |
| 11 | AD | C | 6 | 83 | F | W | 12 | ✓ | ✓ |  |  | ✓ |
| 12 | AD | C | 6 | 85 | F | W | 14.5 |  | ✓ |  |  | ✓ |
| 13 | AD | C | 6 | 84 | M | W | 22 |  |  |  |  | ✓ |
| 14 | AD | C | 5 | 80 | F | W | 19 | ✓ |  |  |  |  |
| 15 | AD | C | 6 | 83 | F | W | 3 |  |  |  | ✓ |  |
| 16 | AD | C | 6 | 75 | M | W | 15 |  |  |  | ✓ |  |
| 17 | AD | C | 5 | 67 | M | W | 6 |  |  |  | ✓ |  |
| 18 | AD | C | 6 | 57 | M | W | 17 |  |  |  | ✓ |  |
| 19 | AD | C | 5 | 79 | F | W | 14 |  |  |  | ✓ |  |
| 20 | AD | C | 6 | 85 | F | W | 17 |  | ✓ | ✓ |  |  |
| 21 | AD | C | 6 | 58 | F | B | 78.5 |  |  | ✓ |  |  |
| 22 | AD | C | 6 | 80 | F | W | 4.5 |  | ✓ | ✓ |  |  |
| 23 | AD | C | 6 | 65 | M | W | 34 |  |  | ✓ |  |  |
| 1 | CON |  | 1 | 80 | F | W | 6 |  |  |  |  | ✓ |
| 2 | CON |  |  | 80 | M | B | 21 | ✓ | ✓ |  |  |  |
| 3 | CON |  |  | 85 | M | B | 6 |  |  |  |  | ✓ |
| 4 | CON | 0 | 2 | 81 | M | W | 20 | ✓ | ✓ |  |  |  |
| 5 | CON | 0 | 0 | 80 | F | W | 8 | ✓ | ✓ |  |  |  |
| 6 | CON | 0 | 3 | 80 | M | W | 22 |  |  |  |  |  |
| 7 | CON | 0 | 3 | 82 | M | W | 14.5 | ✓ | ✓ |  |  | ✓ |
| 8 | CON | 0 | 4 | 86 | M | W | 7 |  | ✓ |  |  |  |
| 9 | CON | 0 | 2 | 87 | F | W | 7 |  |  |  |  | ✓ |
| 10 | CON | 0 | 2 | 85 | F | W | 27 |  |  | ✓ |  | ✓ |
| 11 | CON | 0 | 1 | 64 | M | W | 28 |  |  |  | ✓ |  |
| 12 | CON | 0 | 2 | 68 | F | W | 12 |  |  |  | ✓ |  |
| 13 | CON | 0 | 2 | 94 | M | W | 15 |  |  |  | ✓ |  |
| 14 | CON | 0 | 2 | 79 | F | W | 33 |  |  | ✓ |  |  |
| 15 | CON | 0 | 1 | 86 | M | B | 38.5 |  |  | ✓ |  |  |
| 16 | CON | 0 | 2 | 89 | F | W | 4 |  |  |  |  | ✓ |
| 17 | CON | A | 3 | 90 | F | B | 22 |  |  |  | ✓ |  |
| 18 | CON | 0 | 0 | 41 | M | W | 12 |  |  |  | ✓ |  |
| 19 | CON | 0 | 1 | 56 | F | B | 7 |  |  | ✓ | ✓ |  |
| 20 | CON | 0 | 2 | 88 | F | W | 9 | ✓ | ✓ | ✓ |  | ✓ |
| 21 | CON | A | 3 | 100 | F | W | 11 |  |  | ✓ |  |  |

**Supplementary Table 2.**
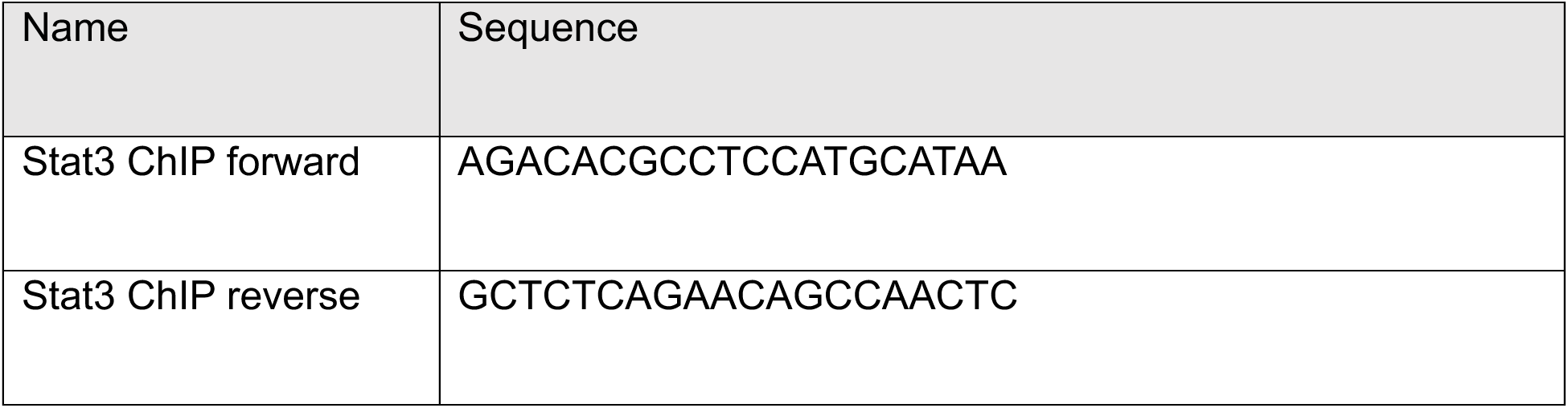
Primers used for ChIP assay from hippocampus of CamK-PARIS mice.

**Supplementary Table 3.**
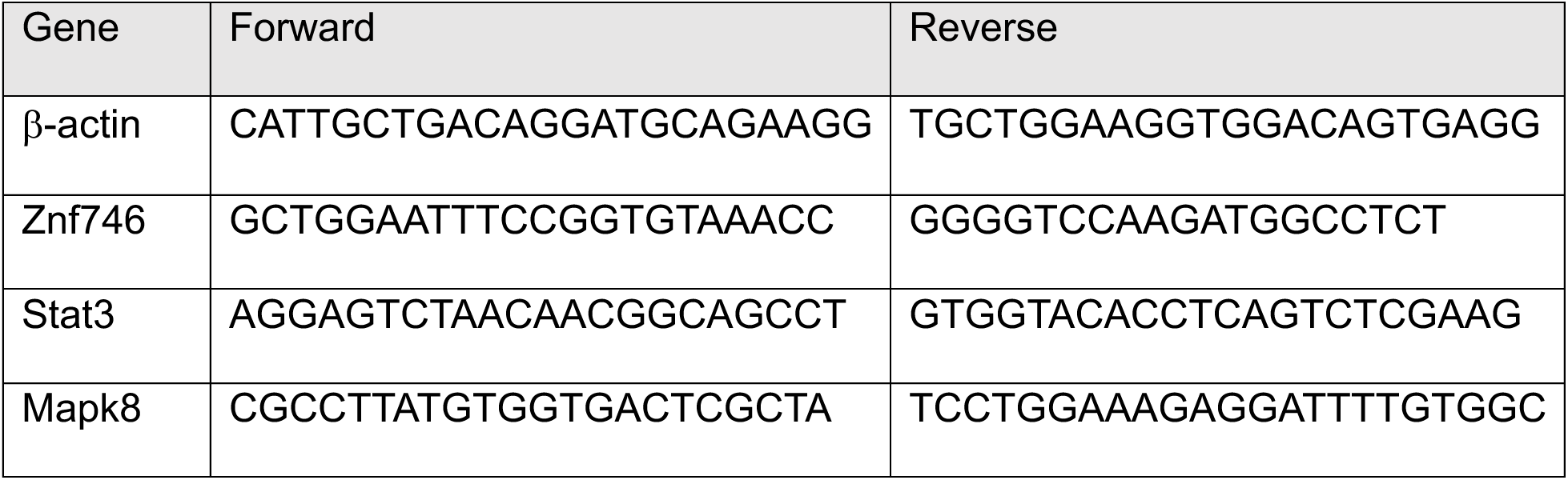
Primers used for qRT-PCR analysis from hippocampus of CamK-PARIS mice.

